# Fluorogenic U-rich internal loop (FLURIL) tagging with bPNA enables intracellular RNA and DNA tracking

**DOI:** 10.1101/2022.07.21.501035

**Authors:** Yufeng Liang, Sydney Willey, Yu-Chieh Chung, Yi-Meng Lo, Shiqin Miao, Sarah Rundell, Li-Chun Tu, Dennis Bong

## Abstract

We introduce herein a new strategy for intracellular RNA and DNA tracking that is robust, orthogonal and complementary to existing methods: <u>F</u>luorogenic <u>U</u>-<u>R</u>ich <u>I</u>nternal <u>L</u>oop (FLURIL) tagging with cell-permeable fluorophore-labeled bifacial Peptide Nucleic Acids (fbPNAs). Our approach uses an 8-nt (U_4_xU_4_) U-rich internal loop (URIL) in the RNA of interest (ROI) as a compact labeling site for fluorogenic triplex hybridization with a bPNA probe (~1 kD). FLURIL tagging thus replaces a 4 bp duplex stem with a labeled 4-base-triple hybrid stem of similar structure and mass. In contrast to existing strategies for RNA tracking, FLURIL tagging can be applied to internal, genetically encoded URIL RNA sites *with minimal structural perturbation*, co-expression of protein-fusion labels or significant increase in molecular weight and steric bulk. We demonstrate effective FLURIL tagging of intracellular (HEK-293) RNAs, ribonucleoprotein (RNP) complexes and live cell (U2-OS) tracking of genomic loci. FLURIL tracking was internally validated by direct comparison with the most widely used live-cell RNA labeling method, MS2-labeling with MCP-HaloTag and Janelia Fluor dyes. In addition, FLURIL-tagging correctly reported on the endogenous RNP in HEK293 cells formed from TAR DNA binding protein 43 (TDP-43-tdTomato) and UG repeat RNA. The FLURIL strategy was also successfully applied to guide RNA (gRNA) in CRISPR-dCas complexes to enable live cell tracking of a low-copy number genomic locus (IDR3), internally benchmarked against MS2/HaloTag labeling of CRISPR-Sirius gRNA targeted to a proximal locus (IDR2). Notably, FLURIL-tagged IDR2 exhibited similar brightness as loci targeted by CRISPR-Sirius gRNA complexes, which bear 8-MS2 hairpins for protein labeling. Together, these experiments show that FLURIL tagging can simply and reliably track intracellular RNA, RNPs, and DNA, with a streamlined molecular footprint relative to other methods. Importantly, these data also indicate that FLURIL tagging is fully compatible with existing labeling methods without crosstalk and may be used to broaden the scope of intracellular RNA and DNA tracking.

Scheme 1.
FLURIL-tagging of RNAs with bPNA probes.
(a) Triplex hybridization of a U-rich internal loop (URIL) with bPNA (blue) via base triple formation between the melamine base (M) and two uracil bases (inset). (b) General schematic of labeling strategy described herein. An RNA of interest is engineered to contain an URIL and expressed within the cell, with a fluorogenic bPNA probe introduced via cell culture media. Successful URIL targeting is reported by an increase in emission (green) and confirmed by a previously established RNA binding protein with a fluorescent protein (red) fusion.

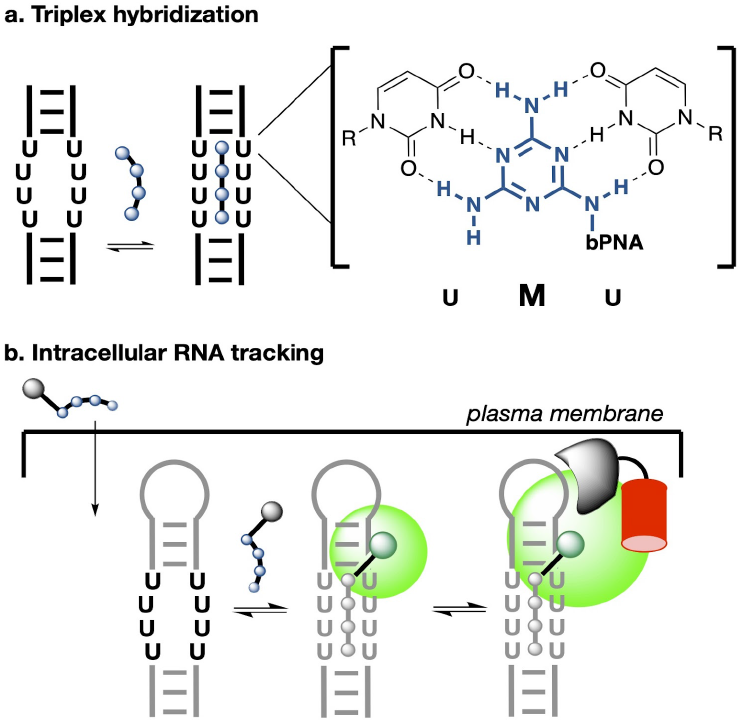

## INTRODUCTION

There remains a need for improved and orthogonal methods for localizing and tracking intracellular RNA and DNA molecules in real-time.^1^ Despite the existence of many useful RNA imaging strategies, there are limitations on placement of probe binding sites and concerns regarding overall structural impact of labeling on the RNA of interest. Modification of internal sites in RNA via chemical reaction or duplex hybridization can be technically challenging or sequence-limited.^2,3^ While fluorescence in situ (duplex) hybridization^4,5^ (FISH) with DNA or PNA probes is broadly used for labeling RNAs, FISH is limited to fixed cell experiments. In addition, duplex hybridization to internal sites destroys existing secondary structures via strand invasion or transforms a single-stranded loop into a new duplex stem where none existed previously,^4^ introducing potentially disruptive steric interactions. Beyond chemical modification and FISH, approaches to intracellular RNA tracking may be roughly divided into small molecule dye binding and protein labels. Dye-binding SELEX-derived aptamers (eg-Spinach,^6^ Broccoli,^7^ Corn,^8^ Mango,^9^ Pepper^10^) have been inserted into RNAs of interest for tracking and sensing applications.^11,12^ A variation on this approach exploits RNA binding to dequench dye-quencher conjugates (eg-SRB2,^13^ Gemini-561,^14^ Riboglow^15^) and can utilize native aptamers to universal quenchers such as cobalamin;^15^ this method may be applied to a wider range of fluorophores without the need for additional aptamer selection. However, all aptamer methods all require insertion of a *distinct folded structure* within the ROI that has the potential to alter native RNA sterics or lifetime. Further, early fluorescent aptamer designs suffered from insufficient brightness thought to be due to misfolded RNA loops.^16^ In particular, the G-quadruplex motif common to Mango and Spinach aptamers^17–19^ is especially labile to intracellular degradation^20^ and requires non-native levels of potassium and magnesium,^7^ which may result in deficient intracellular aptamer performance.^1^ Similarly, bacteriophage-derived RNA hairpins MS2 and PP7 are native sequences that bind to phage coat proteins MCP (<u>M</u>S2 <u>c</u>oat <u>p</u>rotein) and PCP (<u>P</u>P7 <u>c</u>oat <u>p</u>rotein), respectively. The most widely used RNA tracking methodology is to insert MS2 into ROIs and image with MCP fused to fluorescent proteins^21,22^ (MCP-FPs) or MCP-HaloTag^23^ and MCP-SNAP tag^10,24^ fusions, with subsequent dye staining of the MS2 RNPs. Though effective, the use of constitutively fluorescent dyes and FPs generates significant background signal, requiring multiple (often >20) MS2 hairpins to generate sufficient signal to noise. Further, these protein labeling systems carry significant steric bulk (>40 kD for a single MCP-FP) with the risk of altered transcript decay^21,25–27^ and inhibited native contacts. Despite these drawbacks, MS2/PP7 labeling remains the most widely used method for RNA tracking. The use of CRISPR-Cas/guide RNA complexes^28^ for targeting may be used with or without RNA modification, but similarly requires the formation of sterically encumbering synthetic RNPs with native RNA^29^ targets or genomic loci.^30^

Herein, we describe FLURIL-tagging, which utilizes a compact 8-nt (U_4_xU_4_) U-rich internal loop (URIL) that can be selectively labeled with a fluorogenic bPNA probe via triplex hybridization. The FLURIL system minimizes the introduction of new RNA secondary structures by duplex-for-triplex stem replacement and is considerably smaller than a typical aptamer or protein-labeling systems for RNA, yet has comparable brightness and stability. Prior studies on triazine assembly^31–34^ and targeting^35,36^ led to our development of bifacial peptide nucleic acid (bPNA),^33,37,38^ which presents the synthetic melamine base^39,40^ on an α-peptide backbone^41–43^ (Figure 1). The bPNA family of compounds selectively hybridize with U_n_xU_n_ internal bulges^37,38,44,45^ via formation of uracil-melamine-uracil (UMU) base triples, forming in triplex stems that can <u>functionally replace</u> native RNA stems,^44–46^ tertiary contacts,^45,46^ block protein readthrough,^47^ direct chemistry,^48^ and modulate lncRNA lifetime.^49^ Optimized^50^ cationic 4M bPNAs (with 4 melamine bases) are cell-permeable and can bind to structured U_4_xU_4_ bulges while retaining nanomolar affinity,^44^ thus enabling specific intracellular targeting of ROIs that have been engineered to contain a U_4_xU_4_ URIL in the RNA scaffold. This URIL site forms a triplex stem upon hybridization with 4M bPNA that is structurally similar to a 4 bp native duplex RNA stem. Thus, URIL-tagging with bPNA can drive placement of *any sterically-acceptable prosthetic group* into an internal site within the RNA fold without the need for aptamer selection; this manuscript describes the targeting of <u>f</u>luorogen-modified bPNA to the <u>URIL</u> site, which we call <u>FLURIL</u>-tagging. We demonstrate FLURIL-tagging of intracellular RNAs in both fixed and live cell contexts.

**Figure 1.**
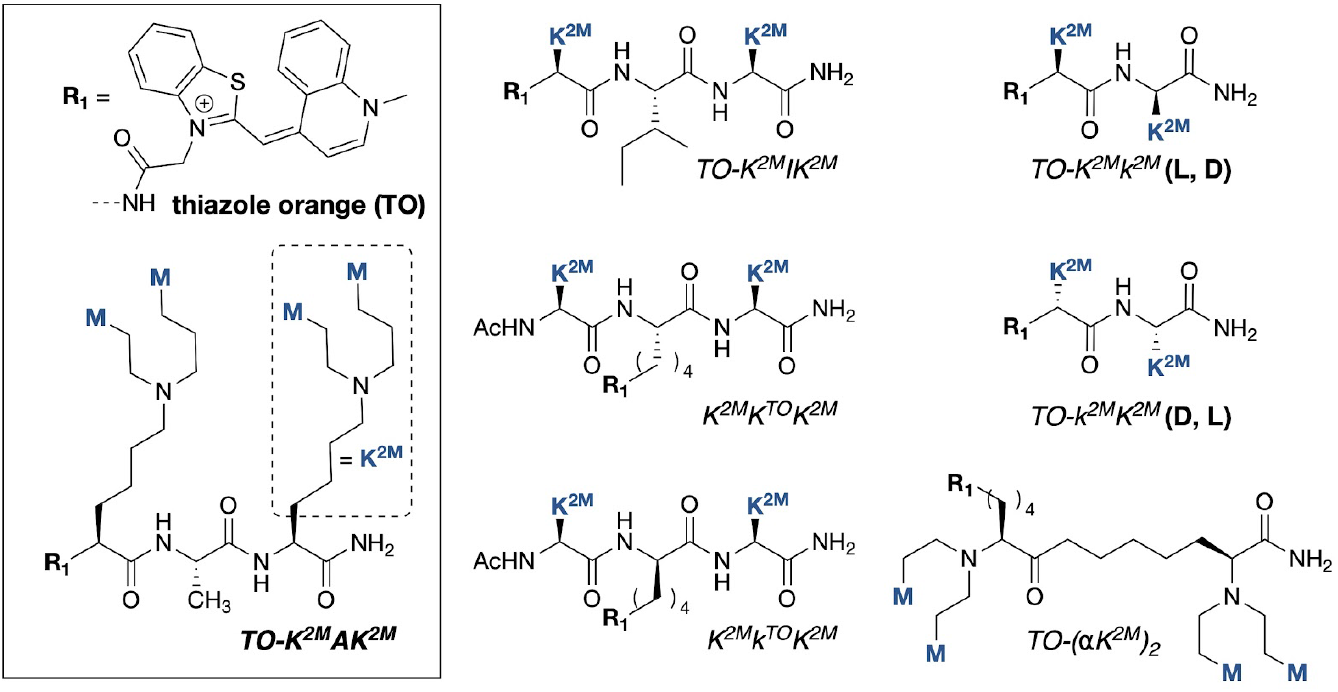
Synthetic bPNA probes. Structures of fluorogenic bPNA probes studied, with thiazole orange (TO) dye and K^2M^ sidechain structure shown inset. Lower case k^2M^ residue denotes D-configuration and K^TO^ indicates sidechain is acylated with R_1_(TO). For TO-K^2M^ AK^2M^ and TO-K^2M^IK^2M^, R_1_ includes a β-alanine spacer between TO and bPNA.

## Results

### Design and synthesis of fluorogenic bPNA probes for URIL RNAs

We previously observed that cyanine dye (Cy3, Cy5) modified bPNAs exhibited fluorescence enhancement upon triplex hybridization to URIL RNA and subsequent hybrid RNA-protein binding,^44^ as presaged^51^ by observations with DNA binding.^52^ Thiazole orange has been more widely used to signal intercalative binding: Seitz and others^53^ have demonstrated sequence-selective “forced intercalation” (FIT) emission with PNA-thiazole orange (TO) conjugates^54^ that place TO at the hybridization interface. Similarly, Dervan has attached TO to pyrrole-imidazole polyamide (Py-Im) minor groove intercalators that fluoresce upon sequence recognition.^55^ While these probes have utility, they are limited with regard to transport and RNA binding.^56^ We synthesized bPNAs *N*-terminated with Cy5, difluorohydroxybenzylidene (DFHBI)^6^ and TO derivatives (Figure 1, Supporting Information). Prior work has established that compact dipeptide or tripeptide bPNA scaffolds containing just 4 melamine base-tripling units (4M bPNAs) can exhibit high affinity binding to a structured U_4_xU_4_ internal bulge. Structure-function studies^50,57,58^ found highly effective nucleic acid binding with the use of a lysine derivative (K^2M^) that presents two melamine bases on the ε-amine; the resulting tertiary amino sidechain becomes cationic upon protonation, obviating the need for additional solubilizing groups and providing electrostatic stabilization of nucleic acid binding. Even following *N*-terminal modification with fluorogenic modules, these efficient bPNA probes are in a low molecular weight regime (~1 kD). Moreover, Fmoc-K^2M^ can be obtained on multigram scale by double reductive alkylation of Fmoc-lysine with melamine acetaldehyde without column purification;^59^ when applied in a di or tripeptide backbone, these bPNA probes are highly accessible and synthetically scalable. We prepared a small family of 4M bPNAs (Figure 1) based on prior structure-function studies, focusing primarily on the tripeptide (K^2M^X-K^2M^) scaffold where X is an α-amino acid, and dipeptide isomers. Within the tripeptide K^2M^Ala-K^2M^ we tested Cy3, Cy5, DFHBI and thiazole orange (TO) dyes at the *N*-terminus. Of these, the turn-on upon RNA complexation was modest for the cyanine and DFHBI dyes (Figures S3.3 & S4.4); we thus focused on TO derivatives. In K^2M^X-K^2M^, thiazole orange carboxylic acid was coupled via a β-alanine linker to the *N*-terminus of (X=Ala, Ile) or directly to the ε-amine of lysine (X=K^TO^). Dipeptide and tripeptide bPNAs of alternating amino acid configuration had previously shown an advantage in triplex hybridization over the homochiral dipeptides due to syndiotactic base presentation^50,60,61^ and we set out to test the impact of stereochemistry on fluorogenic binding when modified with TO on the *N*-terminus (TO-k^2M^K^2M^, TO-K^2M^k^2M^) and ε-amine of the central amino acid (K^2M^k^TO^-K^2M^). Similarly, we synthesized the isodipeptide (TO-(αK^2M^)_2_) that features a sidechain linkage, base presentation on the α-nitrogens and direct TO modification on the ε-amine.

### *In vitro* evaluation of fluorogenic URIL-RNA binding by bPNA variants

With these URIL-RNAs and bPNAs in hand, we tested fluorogenic binding *in vitro* to hairpin and duplex RNAs that were designed to present U-rich internal loops (URILs) in varying contexts (Figure 2A). The absorbance and emission properties of the free TO-bPNAs were in line with the emissively dark thiazole orange parent dye, which has a quantum yield of ~10^-4^ (Supporting Information).^62^ When TO-K^2M^Ala-K^2M^ is bound to URIL RNA (12-U_4_-12, a construct with a U_4_xU_4_ internal bulge buttressed by two 12 bp duplex stems), the hybrid quantum yield (Supporting Information) increases substantively to 43% (Figure 3B), as measured relative to fluorescein (quantum yield=91%). This improvement in TO-bPNA hybrid quantum yield upon URIL binding compares well to the enhancement observed when TO-biotin is bound by the RNA aptamer Mango (quantum yield=14%).^9^ An RNA hairpin construct with a fully base-paired duplex stem (RNAII) does not improve the brightness of the TO-bPNA derivatives (Figure 3C) while binding to URIL-containing RNAs enhanced thiazole orange emission by up to ~600X, with the greatest increase observed between the tripeptide bPNAs and 12-U_4_-12 (Figure 3D). While this degree of fluorescence turn-on upon RNA binding is expected for non-specific intercalation of thiazole orange dyes at higher concentrations,^63^ the null response with the control RNA that lacks a URIL (RNAII) testifies to the essential role of bPNA triplex hybridization in guiding TO binding. All bPNAs gave a strong fluorogenic response to URIL RNAs, but there appeared to be a significant difference in efficacy between the α-linked tripeptides and the isodipeptide bPNA. Further, despite the biophysical advantage previously observed for triplex hybridization with L,D and D,L dipeptide bPNAs, these fluorogenic variants were not as bright as the tripeptides (K^2M^AlaK^2M^, K^2M^IleK^2M^) in this *in vitro* assay, though there may be a slight advantage for K^2M^k^TO^K^2M^over K^2M^K^TO^K^2M^. Based on these results, we chose the TO-K^2M^AlaK^2M^ probe for subsequent intracellular studies.

**Figure 2.**
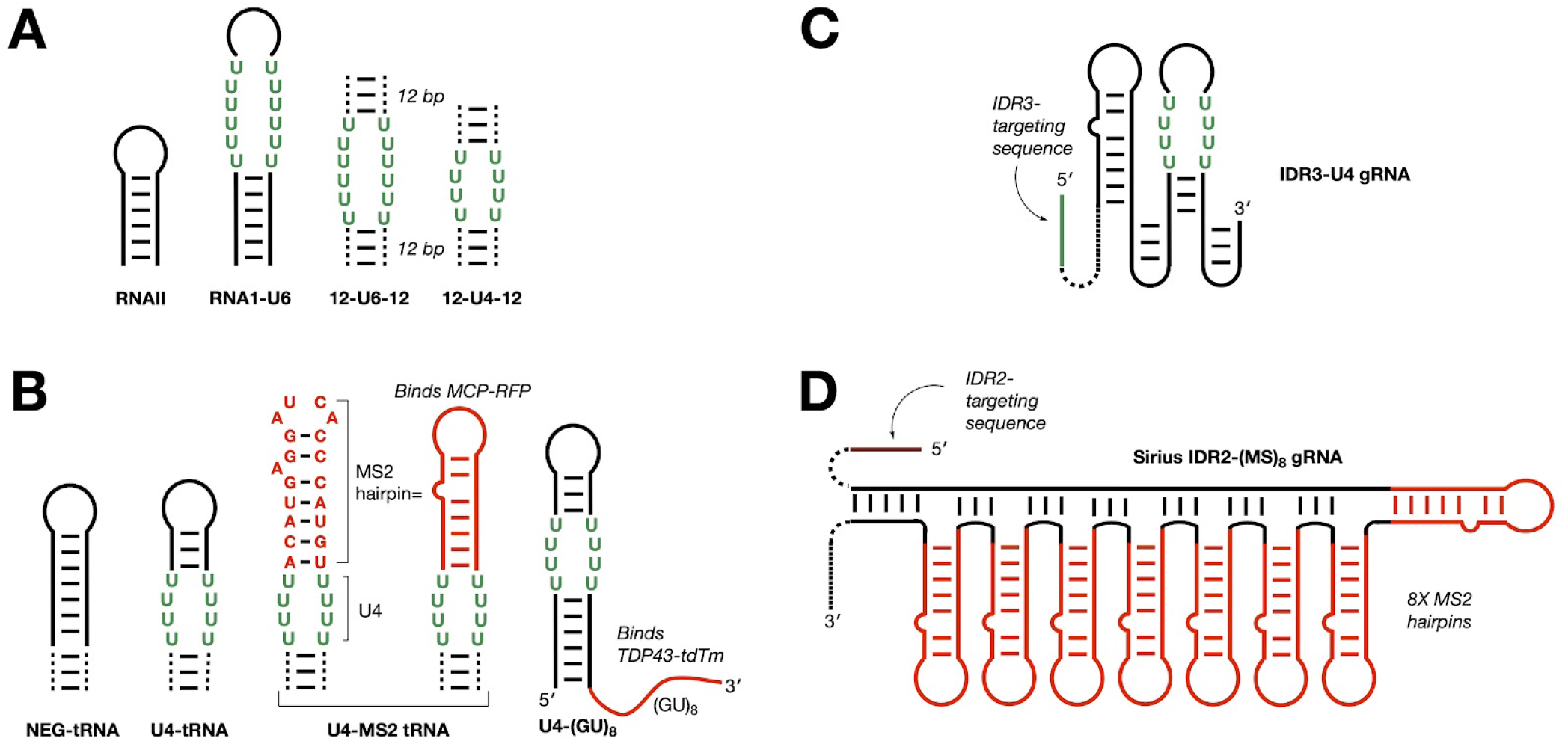
Design of URIL RNA constructs for bPNA probe binding. (A) RNAs for *in vitro* evaluation were prepared via run-off transcription or purchased directly. The 12-U6-12 and 12-U4-12 constructs feature 12 bp duplexes (indicated by dashed lines) separated by U_6_xU_6_ and U_4_xU_4_ internal bulges, respectively. (B) RNA constructs for fixed cell labeling (HEK-293T) of RNPs are shown; hairpin RNAs were used to replace the anticodon stem of tRNA^Lys^ (dashed lines) or appended to a TDP-43 binding (GU)_8_ sequence. Constructs for live cell (U2OS) RNA tracking with (C) FLURIL-tagging and (D) Sirius IDR2-(MS2)_8_ with MS2 hairpins shown in red. IDR3-U4 carries a single PP7 hairpin as an internal control. RNAs for cell experiments were delivered by plasmid transfection. See Supporting Information for full sequence and protocols.

**Figure 3.**
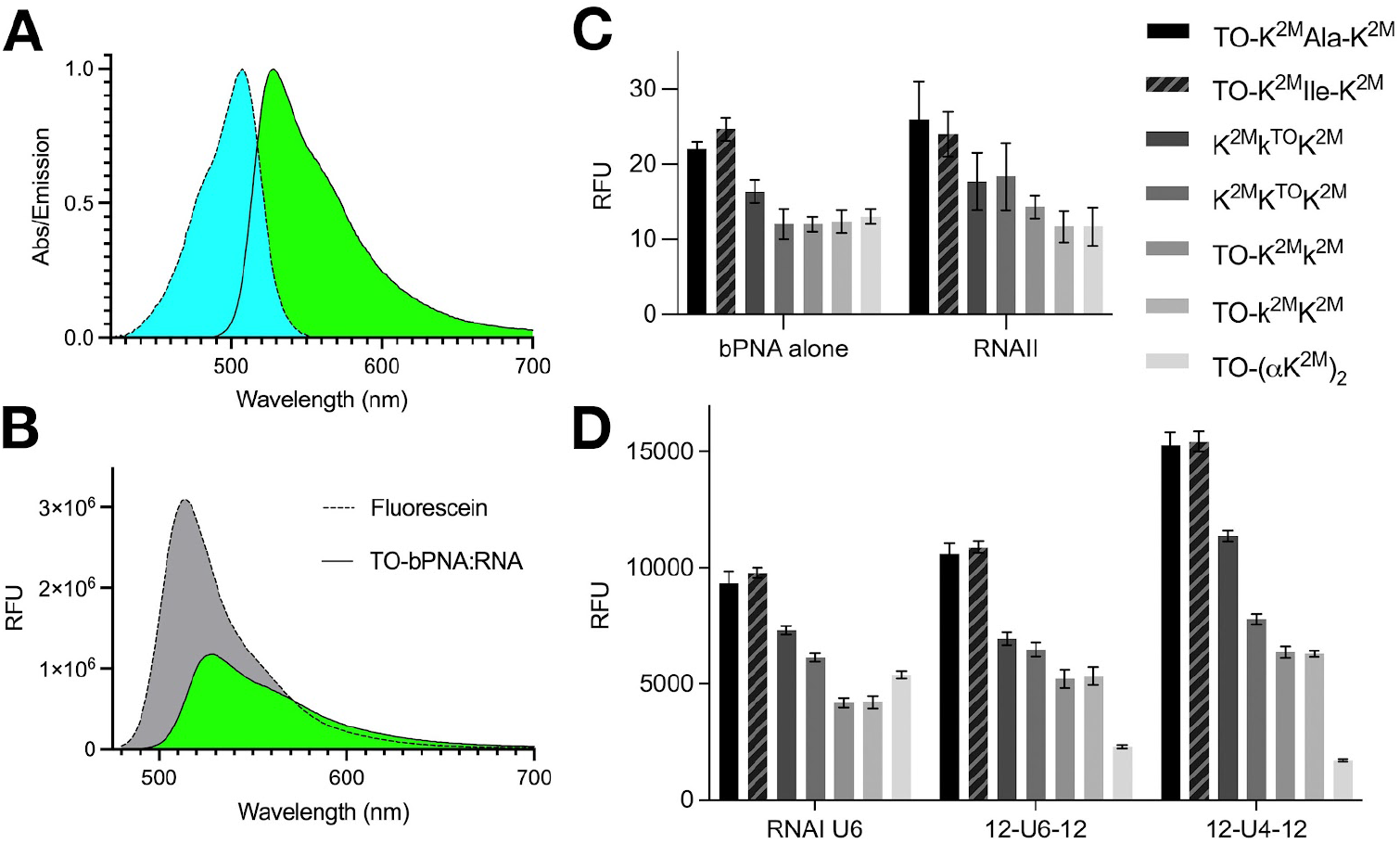
Characterization of fluorogenic bPNA hybridization with URIL RNAs. (A) Normalized absorbance and emission of a representative TO-bPNA (TO-K^2M^Ala-K^2M^) hybrid with RNA. (B) Relative emission of fluorescein standard and the hybrid in (A) indicating relative brightness (Φ=43%, ε_507_=44,883 M^-1^cm^-1^). *In vitro* fluorescence of (C) bPNAs alone or with RNAII that lacks a URIL and (D) upon treatment with RNA constructs that have a URIL binding site (see Figure 2 for predicted secondary structure). Measurements for all RNA and bPNA combinations were carried out in triplicate under identical conditions with standard deviation error shown.

### Intracellular labeling of URIL RNAs and bacteriophage RNPs with fluorogenic bPNA

Using similar design principles as the *in vitro* studies, URIL RNA hairpins (Figure 2B) were engineered to replace the anticodon loop of a tRNA^Lys^ platform that affords stable intracellular RNA expression^6^ upon plasmid transfection into HEK293 cells. In analogy to the RNAII *in vitro* control, a fully base-paired anticodon stem (NEG tRNA) was designed to serve as an intracellular negative control that lacks bPNA binding. A bPNA-binding target was provided by the U4-tRNA construct, which was used to determine intracellular fluorogenic triplex hybridization with TO-bPNA. Indeed, treatment of HEK293 cells with TO-K^2M^Ala-K^2M^ in cell culture media following transfection with either U4 or NEG tRNA resulted in a 5-10 fold brighter fluorescence intensity in the U4-tRNA transfected cells (Figure 4). These data were supportive of intracellular targeting of the engineered URIL RNA, albeit with diminished enhancement relative to *in vitro* conditions.

**Figure 4.**
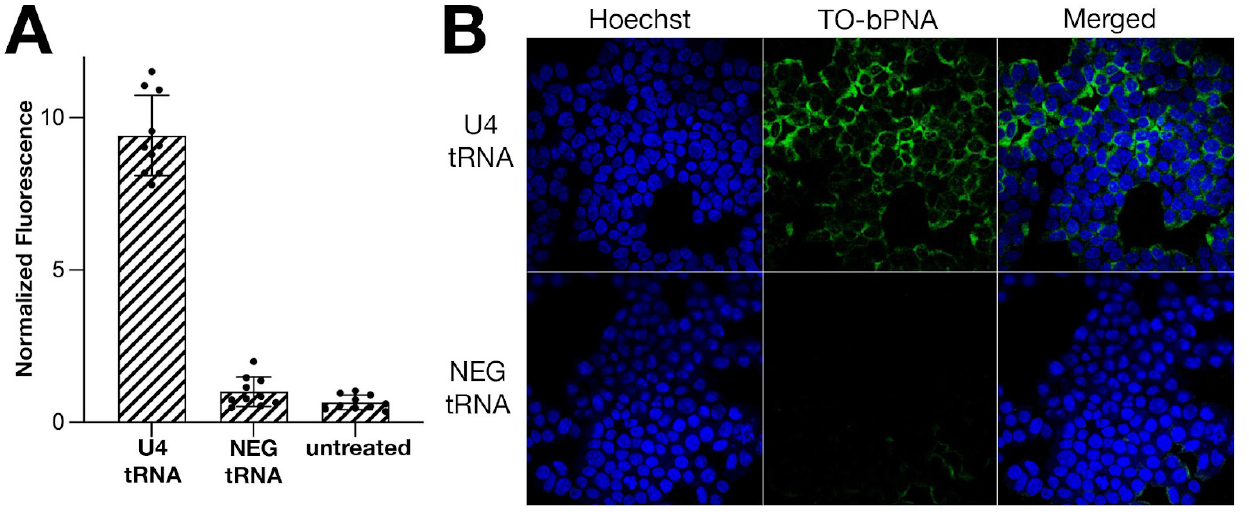
URIL-specific intracellular fluorescence. (A) Integrated fluorescence of the U4-tRNA, NEG-tRNA and untreated cells, normalized to untreated cells. All cell samples were treated with TO-K^2M^Ala-K^2M^. (B) Confocal fluorescence microscopy images of HEK293 cells in the indicated emission channels treated with TO-K^2M^Ala-K^2M^ in media and transfection with plasmid encoding (Top row) U4 tRNA and (Bottom row) NEG tRNA.

To benchmark bPNA labeling of RNA against known RNA tracking strategies, we juxtaposed the U4 URIL with the MS2 hairpin sequence in the tRNA^Lys^ anticodon loop to yield U4-MS2 tRNA (Figure 2B). The U4-MS2 tRNA was co-transfected into HEK293 cells with a plasmid encoding an MCP-RFP fusion, which also bears a nuclear localization signal (NLS).^64^ Cells imaged two hours after transfection and treatment in media with TO-bPNA revealed clear co-localization of red and green fluorescence, supportive of labeling of the U4-MS2 tRNA with TO-bPNA and subsequent binding of the MCP-RFP protein fusion to the triplex hybrid in the cytoplasm (Figure 5). Notably, imaging 8 hours after transfection indicated that both green and red fluorescence had co-localized to the nucleus, consistent with time-dependent transport of both MCP-RFP (which bears the NLS) along with the TO-bPNA hybrid with U4-MS2 tRNA. Thus, treatment of cell culture with TO-bPNA in media effectively labels intracellular URIL RNAs, associated complexes with RNA binding proteins and further, can track intracellular transport of these complexes. Importantly, in the absence of MCP-RFP, green fluorescence from TO-bPNA remains cytoplasmic; in the absence of TO-bPNA, MCP-RFP remains localized to the nucleus as expected.

**Figure 5.**
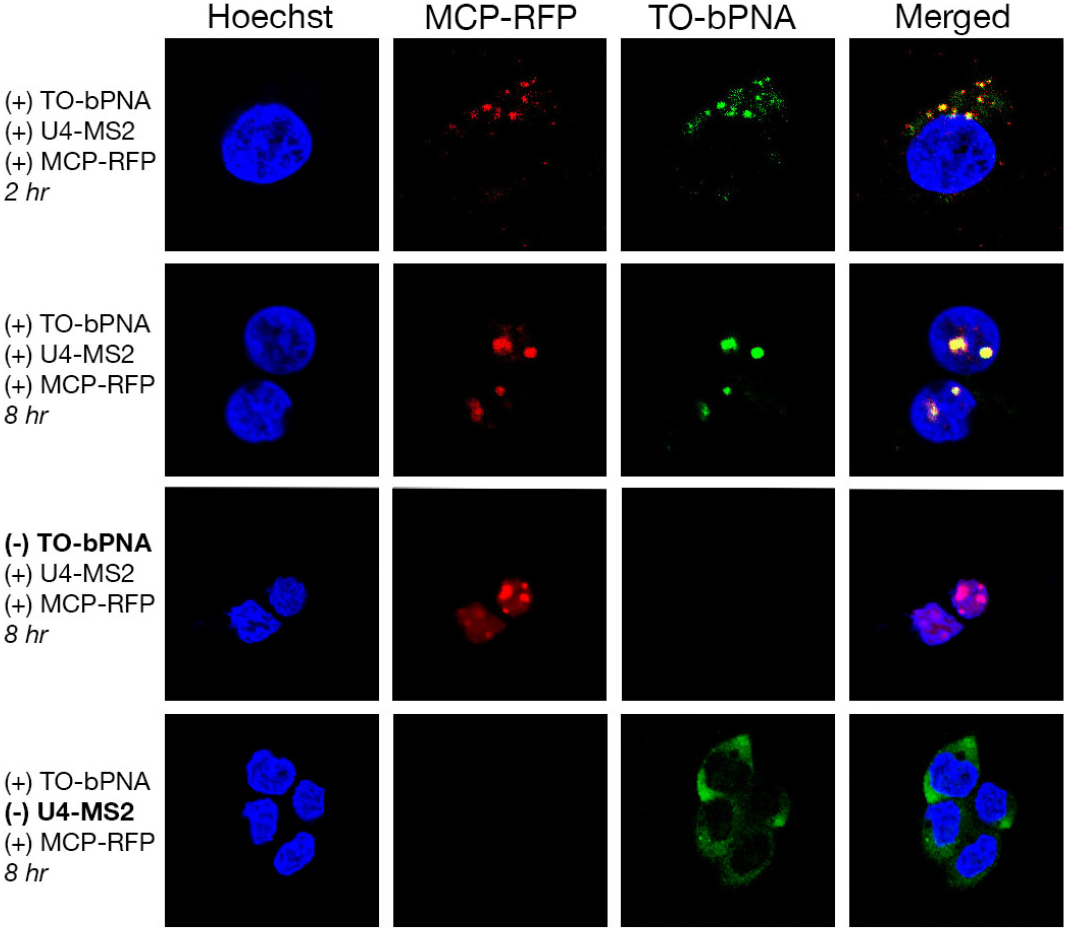
Simultaneous MS2 and FLURIL imaging of RNPs using MCP-RFP and TO-bPNA. Confocal fluorescence microscopy images of HEK293 cells treated as indicated at left of each row and imaged under the dye channels indicated at the top of each column. The MCP-RFP fusion also contains a nuclear localization tag. The first row is imaged two hours after transfection and treatment with TO-K^2M^Ala-K^2M^ (1 μM) in media while subsequent rows are imaged 8 hours after transfection.

### Intracellular labeling of mammalian RNPs with fluorogenic bPNA

To complement the bacteriophage RBP system, we engineered the identical experiment with a native mammalian RNA and protein pair to verify intracellular targeting by TO-bPNA. There have been extensive studies on TAR DNA/RNA binding protein (TDP-43) due to its central role in the progression of neurodegenerative diseases such as ALS,^65–67^ where it is mislocalized from the nucleus to the cytoplasm.^68^ As repeat sequences of UG/TG are established as native RNA partners for TDP-43 repeat sequence,^69,70^ we co-transfected HEK293 cells with plasmids encoding a URIL hairpin pendant to a UG repeat (U4-(GU)_8_, Figure 2B) and TDP-43-tdTomato.^71^ Upon incubation with TO-bPNA in cell media as before, this protocol resulted in the co-localization of red (tdTomato) and green fluorescence in the nucleus (Figure 6). This is again supportive of TO-bPNA reporting on the location of a mammalian RNA-protein complex in the correct subcellular compartment. With the MS2/MCP and (GU)_8_/TDP-43 systems verified, we transfected with swapped RNA-protein pairings that should not result in complex formation: U4-MS2 was co-transfected with TDP-43-tdTomato, and U4-(GU)_8_ with MCP-RFP (Figure 6). In these experiments, both red fluorescent proteins were retained in the nucleus, but the TO-bPNA tagged RNAs remained cytoplasmic, indicating that RNA-protein binding is driving co-localization of red and green fluorescence signals.

**Figure 6.**
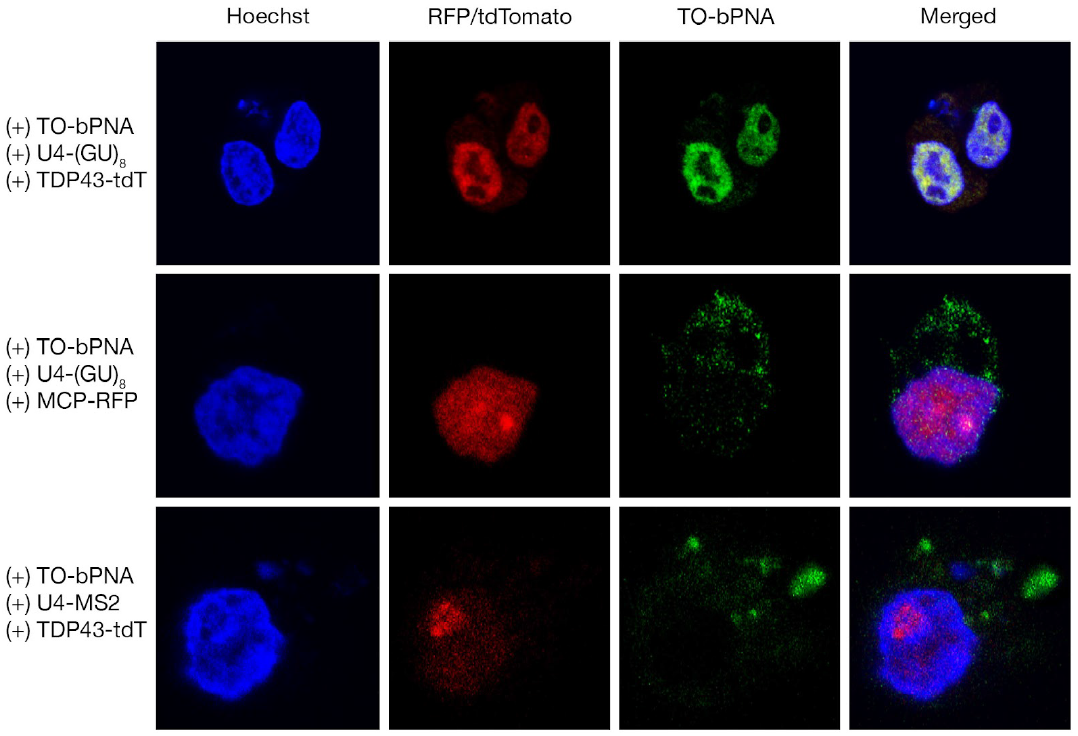
Imaging native RNP pairs using TDP-43-tdTomato and TO-bPNA. Confocal fluorescence microscopy images of HEK293 cells treated as indicated at left of each row and imaged under the dye channels indicated at the top of each column. Top row indicates nuclear colocalization of TDP-43-tdTomato with TO-K^2M^Ala-K^2M^ fluorescence when co-expressed with U4-(GU)_8_. Lower two rows feature mismatches between RNA and RBP (U4-(GU)_8_ with MCP-RFP and U4-MS2 with TDP-43-tdTomato, respectively), resulting in partitioning of TO-K^2M^Ala-K^2M^ emission to the cytoplasm.

### Live cell tracking of genomic loci with bPNA-labeled CRISPR-dCas/gRNA complexes

To test the scope of bPNA cell imaging, we incorporated TO-bPNA reporting into an established approach for live cell genomic loci tracking.^29^ CRISPRainbow is a multiplexed, live cell imaging strategy that uses MS2 or PP7 modified guide RNA (gRNA) complexed to deactivated Cas9 (dCas9) to precisely track endogenous DNA repeats marking specific genomic loci.^72^ Incorporation of MS2/PP7 bacteriophage RNA hairpin sequences into the gRNA enables the dCas-gRNA complexes to be tracked in live cells by fluorescence imaging upon co-expression of phage coat protein (MCP/PCP) fusions with a “rainbow” of fluorescent proteins; display of multiple MS2/PP7 hairpins in a sequence-optimized array yielded more stable and brighter imaging systems known as CRISPR-Sirius gRNAs.^73^ We thus selected two CRISPRainbow/Sirius gRNAs that target proximal intergenic DNA regions IDR2 and IDR3 and modified one to utilize FLURIL-tagging only, with the other serving as an internal benchmark of MS2 labeling. On the gRNA targeting IDR3, one bacteriophage hairpin domain was deleted and replaced with a single U_4_XU_4_ bulge (URIL) in a stabilized hairpin (Figure 2C, Supporting Information), rendering the construct (IDR3-U4) trackable by FLURIL-tagging. The IDR3-U4 gRNA carries a PP7 hairpin that serves as an internal control for labeling experiments that lack bPNA probe, and was indeed found to be unreactive to labeling in the absence of TO-bPNA. To target IDR2, a CRISPR-Sirius gRNA called Sirius-IDR2-(MS2)_8_ was used (Figure 2D), which features an array of 8 MS2 hairpins optimized for stability. A plasmid was constructed encoding both IDR3-U4 and Sirius-IDR2-(MS2)_8_ and this vector was transfected into U2OS cells stably expressing dCas9 and MCP-HaloTag. Staining of live cells in culture with TO-bPNA and HaloTag-JF549 dye thus enabled simultaneous and orthogonal imaging of IDR3 and IDR2, respectively (Figure 7A).

**Figure 7.**
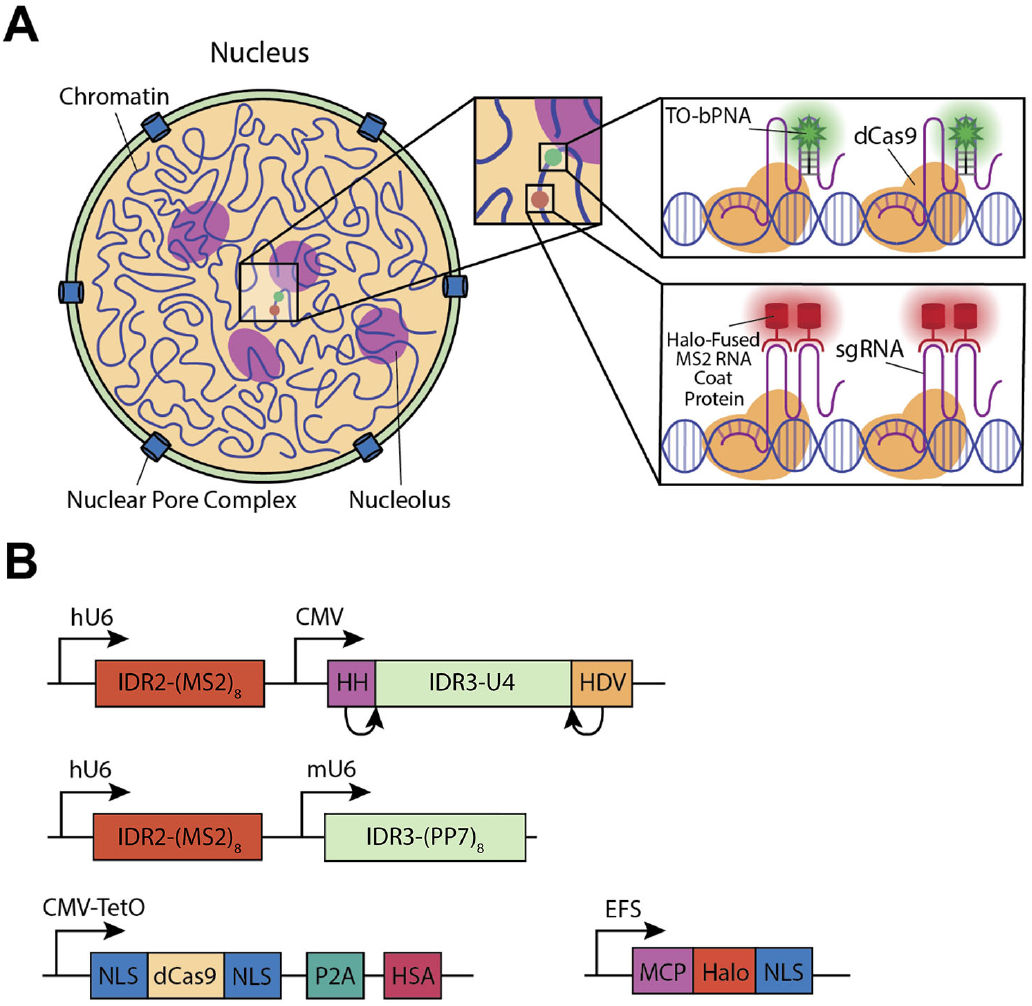
FLURIL-tags in CRISPR-dCas live cell genomic loci tracking. (A) Illustration of dual color genomic labeling of IDR2 and IDR3 by CRISPR-dCas9 targeting. IDR3 is tracked by FLURIL-tagging of IDR3-targeting gRNA while MS2-labeling of IDR2-targeting gRNA is used to track IDR2. FLURIL-tags are stained with TO-bPNA while MS2 labels are stained with MCP-HaloTag binding and HaloTag-JF549 reaction with the complex. MS2 labeling is accomplished using the CRISPR-Sirius gRNA design in Sirius-IDR2-(MS2)_8_ gRNA which has an array of 8 MS2 hairpins. (B) Plasmids used: (Top) dual-gRNA plasmid driven by two promoters (hU6 for Sirius-IDR2-(MS2)_8_, CMV for IDR3-U4), (Middle) control dual gRNA plasmid with IDR3-U4 gRNA replaced by Sirius-IDR3-(PP7)_8_, (Bottom) plasmids carrying dCas9 under an inducible promoter, MCP-Halotag fusion under continuous expression. NLS=nuclear localization signal; P2A=cleavage peptide; HSA=mouse heat stable antigen.

Final vector design was driven by the low efficiency of initial efforts in expression of the modified gRNAs. We speculated that the oligo-U domains triggered early termination by RNA Pol III, which is known to significantly reduce the gRNA targeting efficiency in the CRISPR system.^64,74^ To bypass this issue, we expressed gRNA under a CMV Pol II promoter and ensured precise transcript generation by flanking the gRNA sequence with Hammerhead (HH) and Hepatitis Delta Virus (HDV) self-cleaving ribozymes.^75^ We thus modified the gRNA plasmid to contain the Hammerhead (HH) and Hepatitis delta virus (HDV) ribozymes on either side of the gRNA under a CMV promoter, and inserted it into a dual gRNA plasmid expressing CRISPR-Sirius gRNA (Sirius-IDR2-(MS)_8_) for dual-color genomic loci targeting (Figure 7). Indeed, the resulting dual color genomic loci labeling system for IDR2 (Sirius-IDR2-(MS2)_8_ + MCP-Halotag + Halotag-JF549) and IDR3 (IDR3-U4 + TO-bPNA) was readily established in U2OS cells by lipofectamine plasmid transfection and staining with dyes delivered in media (HaloTag-JF549, TO-bPNA). Live cell imaging revealed partially overlapped red (IDR2, HaloTag-JF549) and green (IDR3, TO-bPNA) foci (Figure 8 top row), which was expected given the close proximity (~4.6 kB) of the two loci on chromosome 19. Exclusion of TO-bPNA from media resulted in loss of IDR3 foci, leaving only red IDR2 foci visible (Figure 8 middle row). Additionally, identical plasmid transfection protocols were carried out in which the FLURIL gRNA was replaced with a CRISPRainbow gRNA bearing PP7 hairpins (Sirius-IDR3-8xPP7). When stained with HaloTag-JF549 and TO-bPNA, only red foci were visible (Figure 8 bottom row), consistent with URIL-selective fluorogenic binding of the TO-bPNA probe previously observed in fixed cells (Figure 4). FLURIL-tagged IDR3 loci was easily tracked over 13s without significant photobleaching or dye blinking, performing comparably to IDR2 loci tracking with the established CRISPR-Sirius (8X-MS2) system.^73^ Time-lapse imaging was carried out for 96 frames at an capture rate of 136 ms per frame (movie S1), yielding IDR2 and IDR3 trajectories on identical and homologous chromosomes of similar range (~100-200 nm) but in different patterns of loci territories, consistent with labeling of proximal, but distinct loci (Figure 9). FLURIL-tagging of gRNA thus gave a highly usable fluorescence signal-to-noise ratio with single locus labeling, likely due to minimal background fluorescence from unbound TO-bPNA, unlike constitutively fluorescent FP and HaloTag systems. Though single molecule fluorescence tracking has not been demonstrated, we note that the sole FLURIL-tag was sufficient for labeling the low-copy number IDR3 locus (45 copies over 1.5 kB). Under standard operating conditions, this single FLURIL gRNA tag affords similar brightness to the CRISPR-Sirius gRNA tags that have a substantially larger molecular footprint of 8 bacteriophage RNA hairpins bound by bacteriophage coat protein fusions.

**Figure 8.**
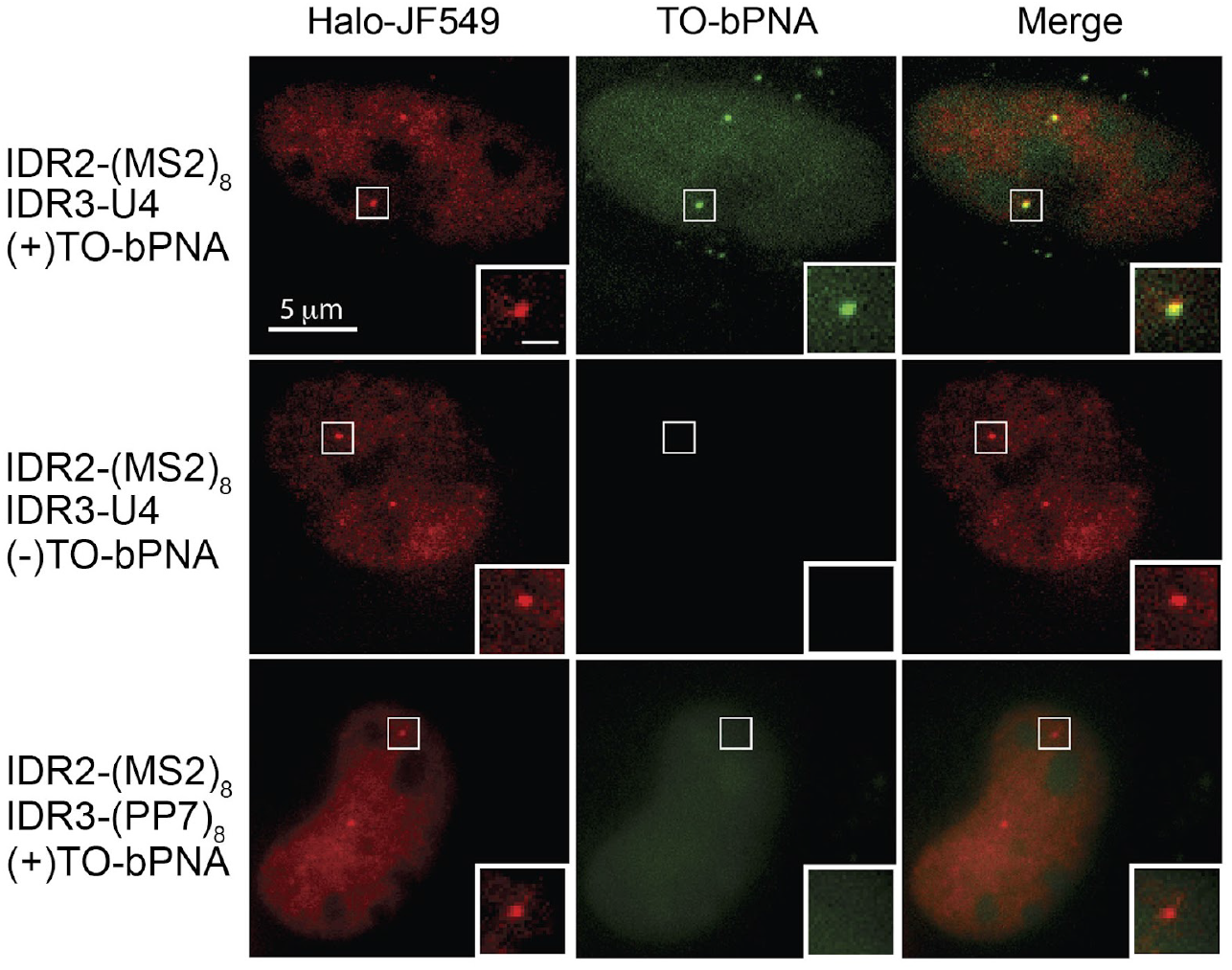
Dual-color CRISPR imaging of IDR2 and IDR3 genomic loci in U2OS by simultaneous FLURIL-tagging and MS2-labeling of gRNAs. (Top Row) Loci imaging following transfection with dual plasmid encoding Sirius-IDR2-(MS2)_8_ and IDR3-U4 followed by staining with Halo-JF549 (red) and TO-bPNA (green), respectively. Genomic loci highlighted in the white square are shown enlarged, inset bottom right-hand corner. (Middle Row) Same plasmid transfection as top row without TO-bPNA treatment. (Bottom Row) Loci imaging following transfection with dual plasmid encoding Sirius-IDR2-(MS2)_8_ and Sirius-IDR3-(PP7)_8_ followed by staining with Halo-JF549 (red) and TO-bPNA (green), respectively. Bar in the inset, 1 μm.

**Figure 9.**
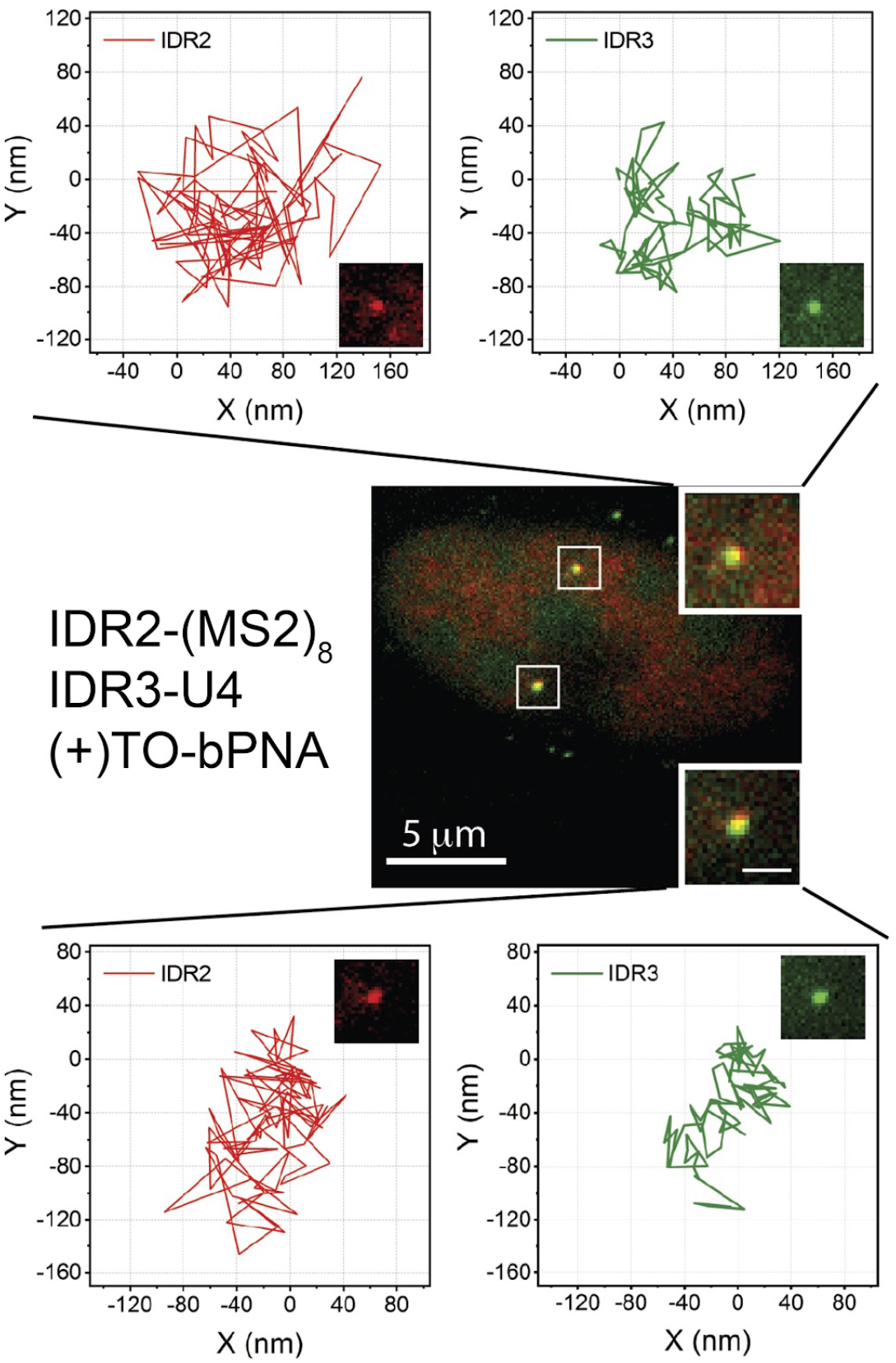
FLURIL-tag Tracking of IDR2 and IDR3 dynamics in live cells. U2OS cells were imaged 48 hours post-transfection with dual plasmid encoding Sirius-IDR2-(MS2)_8_ and IDR3-U4 and stained with HaloTag-JF549 (red) and TO-bPNA (green). Loci trajectories of 96 frames from homologous chromosomes are shown in the top and bottom. The imaging rate is 136 ms per frame. Bar in the inset, 1 μm. For comparison, trajectories were aligned to start from the origin (0,0).

## Discussion

Taken together, these data establish FLURIL-tagging as an efficient and convenient strategy for intracellular RNA, RNP, and DNA tracking that leverages selective intracellular triplex hybridization of cell-permeable bPNA to URILs with fluorogenic binding to establish a robust and bright RNA tracking signal. We find that TO-bPNA exhibits significant (~600X) enhancement of emission intensity upon bPNA triplex hybridization with structured RNAs URILs *in vitro* and a substantive increase in quantum yield from ~0.01% in the unbound state to 43% in the RNA complex. This significant fluorogenic response to URIL-RNAs translated to 5-10X intracellular signal-to-noise and a highly usable, low-background cellular imaging of URILs. This method requires a modest 4 bp stem site within an RNA of interest to install an 8 nt URIL and plasmid transfection to introduce the modified RNA. While an engineered biomolecule (URIL-RNA) is needed, this is a universal requirement for live cell tracking strategies. Intracellular FLURIL-tagging of RNAs in both fixed and live cells was directly verified using established protein-labels from endogenous mammalian RNPs as well as bacteriophage MS2-labeling, which remains the gold standard for RNA tracking. Indeed, RBP fusions with fluorescent proteins (FP) confirmed that FLURIL-tags were labeling the correct RNA species, while cells lacking the correct RNA-protein pairing resulted in segregation of FLURIL tag and RBP-FP fusion signals. Under live cell tracking conditions, FLURIL tags also performed comparably to established MS2-based methods (CRISPR-Sirius), while occupying a considerably more compact molecular footprint. It is particularly noteworthy that the FLURIL tag only requires replacement of a 4 bp stem with an 8 nt URIL to enable RNP live cell tracking with a cell-permeable, ~1kD bPNA probe. In the current study, we have inserted stabilized hairpin structures to host the URIL site and included a PP7 hairpin as a control element in IDR3-U4; however, a URIL could be inserted into a native fold by replacement of an existing native 4 bp stem buttressed by other structures in the ROI, further streamlining the FLURIL-tag footprint. Though there remain many appealing aspects to MS2 labeling including multiplex and multicolor approaches, the use of protein fusions and multiple RNA hairpin sites encumbers the RNA of interest with substantial steric bulk, adding 41-47 kD of mass per MCP-FP and MCP-HaloTag fusion, respectively. The steric bulk and RNA secondary structures required for protein labeling have raised concerns that this could affect native transcript processing and tertiary contacts;^1^ however, inhibition of transcript degradation by insertion of bacteriophage (MS2/PP7) hairpins is less of an issue in mammalian cells.^21,26,27^ In contrast, FLURIL tags add negligible additional mass to the RNA target and a structural perturbation which may be as minor as replacement of a duplex stem with a triplex stem. Prior work has demonstrated that this replacement does not disrupt proximal RNA domains, retaining tertiary interactions as well as catalytic function. The FLURIL tag also avoids the use of labile^20^ G-quadruplex domains and insertion of foreign secondary structures found in some SELEX-derived dye-binding aptamers such as Spinach and Mango. Furthermore, as fluorogenic dye binding is driven by bPNA triplex hybridization to the URIL rather than direct RNA recognition of the dye, new prosthetic groups^49^ may be used to decorate RNPs without the technical demand of a SELEX campaign. In particular, other fluorogens in the cyanine family could potentially be used in FLURIL tagging, thus expanding the color range available without laborious aptamer selection procedures. Analogously, the aptamer Pepper^10^ binds a family of fluorogen dyes with significant structural variation, suggesting that minimal specific contacts are needed to access fluorogenic binding. Efforts to expand both color selection and sequence scope^40^ for FLURIL tagging are currently underway. The current work demonstrates the facility of the FLURIL tag, as well as an appealing compatibility with popular methods such as MS2-labeling. The combination of *orthogonality and compatibility* of FLURIL tags to existing and widely-used technologies that include MS2/PP7, HaloTag, dye aptamers, Riboglow, dCas13-gRNA, FISH and other indicates that RNA FLURIL tagging is a convenient and enabling tool for discovery that broadens the versatility of RNA tracking platforms.

## Supporting information

Movie S1 live cell tracking

## Acknowledgement

We thank Andy Sak and Bern Kohler for guidance on quantum yield determination. We thank the Department of Biological Chemistry and Pharmacology at OSU Medical Center for use of the BD FACSAria Fusion sorter. This work was supported by the NIH 1R01GM143543-01 (D.B.), NIH 4R00GM126810-05 (L.T.), P30 CA016058 (Ohio State University Comprehensive Cancer Center), Ohio State University start up funds (L.T.), STEP fellowship (S.W.) and the Center for RNA Biology at the Ohio State University.

