## Supporting Information for "Fluorogenic U-rich internal loop (FLURIL) tagging with bPNA enables intracellular RNA and DNA tracking"

#### Table of contents


| Section | | Page |
|---|---|---|
| **S1** | Materials and handling | 2 |
| **S2** | General experimental procedures | 4 |
| **S3** | Additional fluorescence microscopy data (fixed cells) | 7 |
| **S4** | Additional in vitro fluorogenic binding data | 9 |
| **S5** | Synthetic procedures | 10 |
| **S6** | Compound characterization | 14 |

**S1. Materials and handling**
All chemicals were used without further purification from commercial sources as indicated, unless otherwise noted. DNAs and RNAs were purchased from Integrated DNA Technologies (IDT). Nucleic acid strands shorter than 20 nt were used without further purification. Otherwise, the DNAs/RNAs were purified by TBE-urea denaturing gel. SYBR® gold was purchased from Thermo Fisher Scientific. DNA stock solutions were serially diluted in MilliQ water and concentrations were determined by measuring solution absorbance at 260 nm on a Thermo Fisher Nanodrop 2000. Sample fluorescence was measured on a Thermo Fisher Nanodrop 3300.

**S1.1. Nucleic acid sequences**

- RNAs for *in Vitro* fluorescence turn-on

A and B strands were annealed together into duplexes before use.
12-U4-12 A: 5’-CGCAUAGCUCAGUUUUGACUCGAUACGC-3’
12-U4-12 B: 5’-GCGUAUCGAGUCUUUUCUGAGCUAUGCG-3’

12-U6-12 A: 5’-CGCAUAGCUCAGUUUUUUGACUCGAUACGC-3’
12-U6-12 B: 5’-GCGUAUCGAGUCUUUUUUCUGAGCUAUGCG-3’

RNAI U6: 5’-GGCAGCUUUUUUUUGGUAGUUUUUUGCUGCC-3’

RNAII WT: 5’-GCACCGCUACCAACGGUGC-3’

- Plasmid delivered RNA sequences for intracelluar labeling

U4 tRNA:
5’-GCCCGGAUAGCUCAGUCGGUAGAGCAGCGGCCGUUUUCGCUCCGGCGUUUUCGGCCGCGGGUCCAGGGUUCAAGUCCCUGUUCGGGCGCCA-3’

U4-MS2 tRNA:
5’-GCCCGGAUAGCUCAGUCGGUAGAGCAGCGGCCGUUUUCGCACAUGAGGAUCACCCAUGUGCGUUUUCGGCCGCGGGUCCAGGGUUCAAGUCCCUGUUCGGGCGCCA-3’

Negative tRNA:
5’-GCCCGGAUAGCUCAGUCGGUAGAGCAGCGGCCGCGCGCGCUCCGGCGCGCGCGGCCGCGGGUCCAGGGUUCAAGUCCCUGUUCGGGCGCCA

$(GU)_8$-U4 RNA: 5’-CGGCCGUUUUCGCUCCGGCGUUUUCGGCCGGUGUGUGUGUGUGUGU-3’

Predicted secondary structures based on previously reported parameters and using ViennaRNA package are shown below in Table S1.[1–3]

**Table S1.** Predicted RNA secondary structures for URIL RNA probe constructs.[1–3]

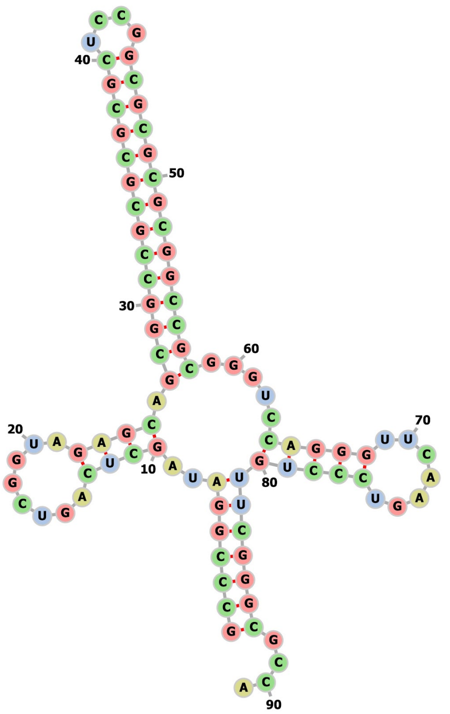

**NEG tRNA** predicted fold.

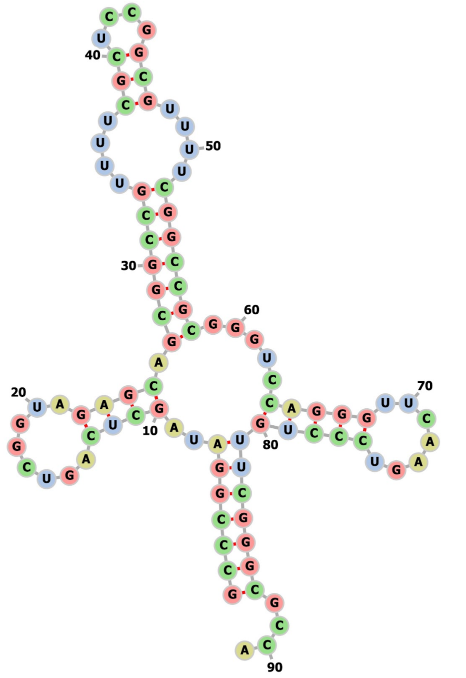

**U4-tRNA** predicted fold.

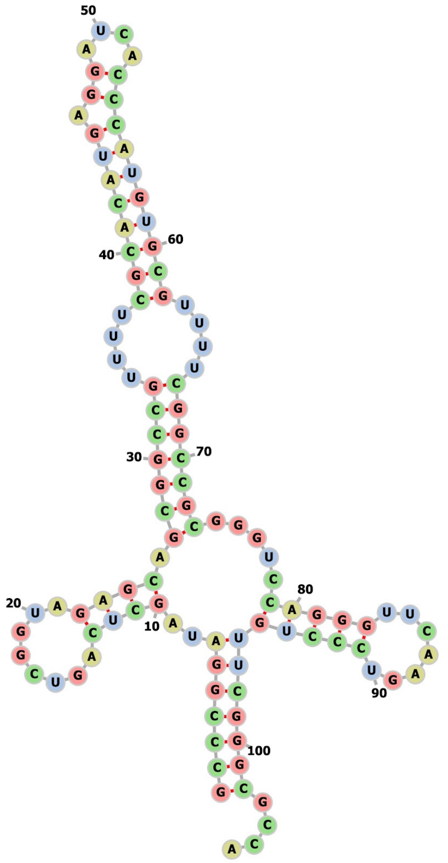

**MS2-U4** tRNA predicted fold.

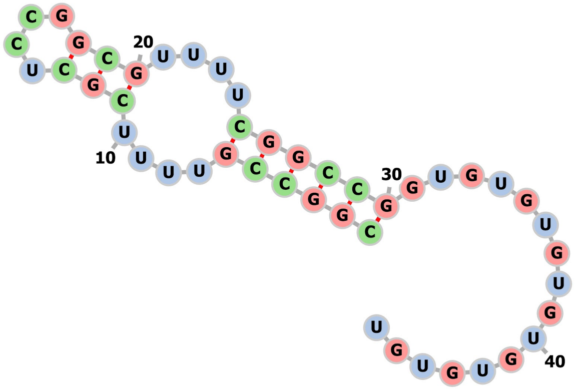

**U4-(GU)$_8$** predicted fold.

### S2. General experimental procedures

#### S2.1. HPLC

Fluorogenic bPNA probe was purified using HPLC: Hitachi D-7000 (interface), Hitachi L-7150 (UV-detector) and Hitachi L-7400 (pump). Analytical and semi-preparative HPLC was carried out with $C_{18}$ reverse phase columns. HPLC solvent A: 99% MilliQ $H_2O$, 1% HPLC grade acetonitrile, 0.1% trifluoroacetic acid; HPLC solvent B: 10% MilliQ $H_2O$, 90% HPLC grade acetonitrile, 0.07% trifluoroacetic acid.

#### S2.2. In vitro fluorescence turn-on measurement

Samples were prepared as follows: 50 mM HEPES (pH 7.5), 100 mM NaCl, 2 μM RNA, 2 μM probe. The nucleic acids were annealed from 95 °C to pre-form the structure before use. The samples were prepared and incubated at room temperature for 30 min before measuring the fluorescence (RFU) with Thermo Fisher Nanodrop 3300 (excitation=470 nm, emission=522 nm). For each measurement, 2 μL sample volume was applied to the center of the nanodrop probe. All fluorescence values are the average of triplicate measurements, starting from fresh sample preparation. Error bars indicate standard deviation.

#### S2.3. Fitting methodology for binding curves

Probes (bPNA-TO) were held at a constant concentration of 100 nM and treated with RNA to final concentrations from 0 to 1.4 μM in buffer (50 mM HEPES, pH 7.5, 100 mM NaCl). The experiments were triplicated from sample preparation and error bars indicate standard deviation. The data were fit with the following equation to obtain dissociation constant Kd :

$$RFU = \frac{m*(Kd+x+100)-m*sqrt[sqr(Kd+x+100)-4*x*100]}{2}$$

#### S2.4. Quantum yield measurement

Fluorescence measurements were taken to obtain relative quantum yield using fluorescein as a standard, which has similar spectral properties to thiazole orange and a known quantum yield of 92% under the experimental conditions.[4] All samples were prepared in 50 mM HEPES (pH 7.5), 100 mM NaCl. RNA used was 12-U4-12 duplex (S1.1). Hybrid samples contained 1 μM RNA, 1 μM TO-bPNA probe; fluorogenic TO-bPNA and free thiazole orange without RNA samples were prepared at 1μM concentration as well. The RNA samples were annealed by slow cooling from 95°C to pre-form the structure before use. The samples were prepared and incubated at room temperature for 30 min before measuring the absorbance with Cary Series UV-Vis-NIR Spectrophotometer. The standard fluorescein was prepared in 0.1M NaOH. All sample concentrations were adjusted to obtain matched sample absorbances, not to exceed 0.04 at 470 nm (± 0.0002). The fluorescence was measured by Quantamaster 8000 spectrofluorometer and corrected for anisotropy with a vertical polarizer in the excitation path (470 nm) and a polarizer in the emission path (507 nm) set at the magic angle (54.7°), with excitation and emission slit widths = 5 nm).[5] Emission spectra were collected and integrated, indicating emission from the bPNA hybrid to be 49.7% that of fluorescein and therefore a relative quantum yield of 43%.

#### S2.5. Construction of RNA plasmid vector

The DNAs were annealed into duplex and from which 2 μg were digested with SalI and XbaI (Thermofisher) in 2x Tango buffer for 12 hours, following protocol provided by Thermofisher. The vector pAV U6+27 (1 μg) was also digested with SalI and XbaI for 12 hours. The digested products were purified on 1% agarose gel and the bands were cut for DNA isolation by QIAquick Gel Extraction Kit (Qiagen). The purified products were ligated (molar ratio of DNA insert:linearized vector=5:1) with T4 DNA ligase (Thermofisher), following the protocol for Corn Aptamer construction.[6] The ligation product was transformed into DH5α for amplification and the transformation mixture was inoculated on Agar plate (Ampicillin). After 16 hours, several colonies were picked and amplified in LB media containing Ampicillin, and the plasmid was isolated by miniprep (Qiagen). The plasmid was sent for sequencing to verify the insertion. Tomato-TDP43 and MCP-TagRFPt plasmids were obtained from Addgene (#28205 and #64541),[7,8] and amplified in the corresponding E.coli cell lines.

#### S2.6. Cell treatment (fixed cell analysis)

HEK-293T cells were cultured based on ATCC protocol. Cells were seeded to 35 mm culture dish (Thermofisher) with clean coverslip (Thermofisher) attached to the bottom at the concentration of $5x10^5$/ml. After one day of incubation, the cells were transfected with corresponding plasmids with Lipofectamine 3000

(Invitrogen) and the cells were incubated for 24 hr. Then the culture medium was removed and the fresh medium containing 1 µM bPNA-dye was added to the attached cells. The cells were incubated at 37°C for 2-8 hr and the medium was discarded. Hoechst 33258 (Invitrogen) in culture medium was added to the cells and incubated for 15 min at 37°C and the medium was discarded. The cells were washed with PBS, fixed with 4% formaldehyde in PBS for 15 min at room temperature and rinsed with PBS. The coverslip was transferred to glass slides for fluorescence microscopy. Cells were imaged under Olympus FV3000 systems. Three or more replicate data sets were obtained that showed consistent findings, starting from cell seeding.

**S2.7. Cellular fluorescence quantification (fixed cell)**
Using ImageJ software, the fluorescence intensity in a circular area of diameter ~5 µm within a single cell (HEK-293T) was selected and calculated. For each treatment, 10 areas were selected in this manner from 10 different cells. The intensity from NEG-tRNA plasmid treated cells was normalized to 1.0.

**S2.8. Plasmid Construction**

The dCas9 (Addgene#121936) and MCP-HaloTag (Addgene#121937) plasmids from prior work in Tu lab were used without modification[9] and expression vector for the guide RNAs (gRNAs) were modified from previous gRNA plasmids. The single gRNA plasmid for IDR3-U4 gRNA was constructed from pLH-sgRNA1-2XPP7[10] (Addgene#75390) where PP7-U4 was inserted to replace 2XPP7 (Figure S1). The dual gRNA plasmid[9] was constructed from pPUR-hU6-Sirius-8XMS2-mU6-Sirius-8XPP7 (Addgene#121944), in which CMV-HH-PP7-4USL-HDV was inserted to replace mU6-Sirius-8XPP7, resulting in pPUR-hU6-Sirius-8XMS2-CMV-HH-PP7-4USL-HDV. The HH and HDV cassettes were inserted for precise gRNA sequences through a cleavage mechanism at a defined position expressed under a pol II promoter in eukaryotic cells. The targeting sequences for IDR2 and IDR3 were 5'-GGAGAGGCTGGG-3' and 5'-AGCAGATGTAGG-3', respectively.[9,11] The gRNA sequences were ordered from IDT and inserted using BbsI restriction enzyme following the same protocol. The modified guide RNA expression vector reported here will be deposited in Addgene.

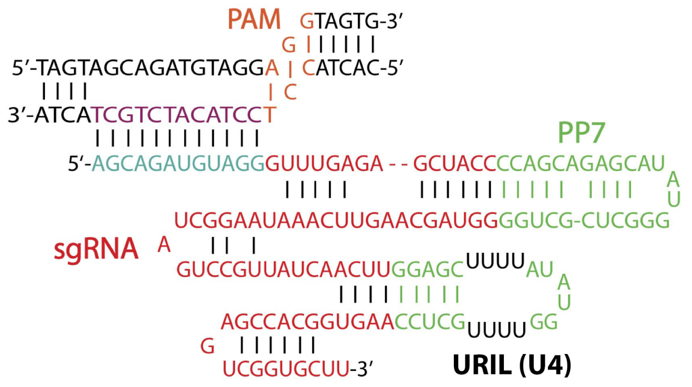


**Figure S1. Sequence and structure of IDR3-U4 single gRNAs.** CRISPRainbow with two PP7 hairpins (2xPP7) gRNA was modified to carry the $U_4xU_4$ internal bulge (URIL) at the second loop position. The cyan color shows crRNA that targets the endogenous DNA sequences (purple). PAM sequence is shown in orange. Red indicates the gRNA scaffold and PP7 and URIL hairpin scaffolds are shown in green. The PP7 hairpin was carried over from a prior sgRNA construct that was modified and serves as an internal control for labeling experiments that lack bPNA probe.

**S2.9. Cell Culture, Lentivirus Production and Transduction (live cell experiments)**
We cultured human osteosarcoma U2OS at 37°C in Dulbecco-modified Eagle's Minimum Essential Medium (DMEM) containing high glucose and supplemented with 10% (vol/vol) fetal bovine serum. $U2OS^{dCas9\text{-}HSA/MCP\text{-}HaloTag}$ cell line that was generated by lentiviral transduction and sorted by FACSFusion cell sorter (BD Bioscience, see S2.10 Flow Cytometry). Lentiviral particles carrying dCas9 and MCP-HaloTag were generated by HEK-293T cells using the same protocol.[11] For lentiviral transduction, U2OS cells maintained as described above were transduced by Spinfection in 6-well plates with lentiviral supernatant for 48 hours and ~$2x10^5$ cells were combined with 1 ml lentiviral supernatant and centrifuged for 30 minutes at 1200 x *g*. For imaging, $U2OS^{dCas9\text{-}HSA/MCP\text{-}HaloTag}$ cells were grown on 35-mm glass-bottom dishes (MatTek) and 2 µg of sgRNA plasmid were transfected using TransIT transfection reagent (Mirus) following manufacturer's protocol. Cells were washed with fresh media 24 hours post-transfection and imaged after another 24 h incubation. A final concentration of 1 µM TO-bPNA was added to the culture medium two hours before imaging. Cell toxicity was

not observed even with prolonged incubation (up to 48 hours) in the presence of TO-bPNA. Cells were tested and found to be free of mycoplasma contamination.

**S2.10. Flow cytometry**

The cell line U2OS$^{dCas9\text{-}HSA/MCP\text{-}HaloTag}$ was generated using the same protocol[9] that generated U2OS$^{dCas9\text{-}HAS/PCP\text{-}GFP/MCP\text{-}HaloTag}$ with the following modifications: (1) The PCP-GFP for labeling PP7 stem loop was not added; (2) cells expressing the dCas9-p2A-HSA and MCP-HaloTag (stained with HaloTag-JF549) were selected using a BD FACSAria Fusion cell sorter (BD Bioscience) equipped with 405, 488, 561 and 640 nm excitation lasers and standard emission filters for PE (582/15) and APC (670/30); (3) AlexaFluor 647-conjugated anti-mouse CD24 antibody (BioLegend) was used to stain for HSA (heat stable antigen, mouse) carried on the dCas9 plasmid; (4) U2OS$^{dCas9\text{-}HSA/MCP\text{-}HaloTag}$ was not selected from a single cell. FACS sorting for dCas9 positive cells was carried out following sample staining (1 µL Alexa Fluor-647 conjugated anti-mouse CD24 antibody, 100 µL cell solution, 30 min). FACS sorting of MCP-HaloTag positive cells was carried out after staining with HaloTag-JF549 (2 nM dye, 12-24 hr).

**S2.11. Fluorescence Microscopy (live cell imaging)**

Cell imaging was carried out on an Olympus IX83 microscope equipped with three EMCCD cameras (Andor iXon 897) mounted on a 4-camera splitter, four lasers (405 nm, 488 nm, 561 nm, and 647nm), mounted with a 1.6x magnification adapter and 60x apochromatic oil objective lens (NA 1.5), resulting in a total of 96x magnification. The microscope stage incubation chamber was maintained at 37°C with $CO_2$ and humidity supplement. A laser quad-band filter set for TIRF (emission filters at 445/58, 525/50, 595/44, 706/95) was used to collect fluorescence signals simultaneously. Data acquisition was carried out with CellSens software. Localization precision was ~5 nm in 4 seconds, ~ 6 nm in 16 seconds, and ~ 10 nm in 80 seconds.[12] The video was recorded 136 ms per frame with a total of 96 frames and 100 ms exposure time. Image size was adjusted to show individual nuclei and intensity thresholds were set on the basis of the ratios between nuclear foci signals to background nucleoplasmic fluorescence.

**Imaging Processing**

The images were registered and analyzed by *Fiji*[13] and *Mathematica* (Wolfram) software. To achieve subpixel registration accuracy, parameters for shifting, scaling, and rotating camera images were determined by the least-squares fitting of fluorescent bead images (100 nm TetraSpeck fluorescent microspheres, Invitrogen). The experimental data from each channel were processed through an affine transformation and overlapped in false-color channels for visualization. The locus trajectory was obtained by the tracking of locus position over time and graphs were generated by *OriginPro* (OriginLab version 2019b).

### S3. Additional fluorescence microscopy data (fixed cells)

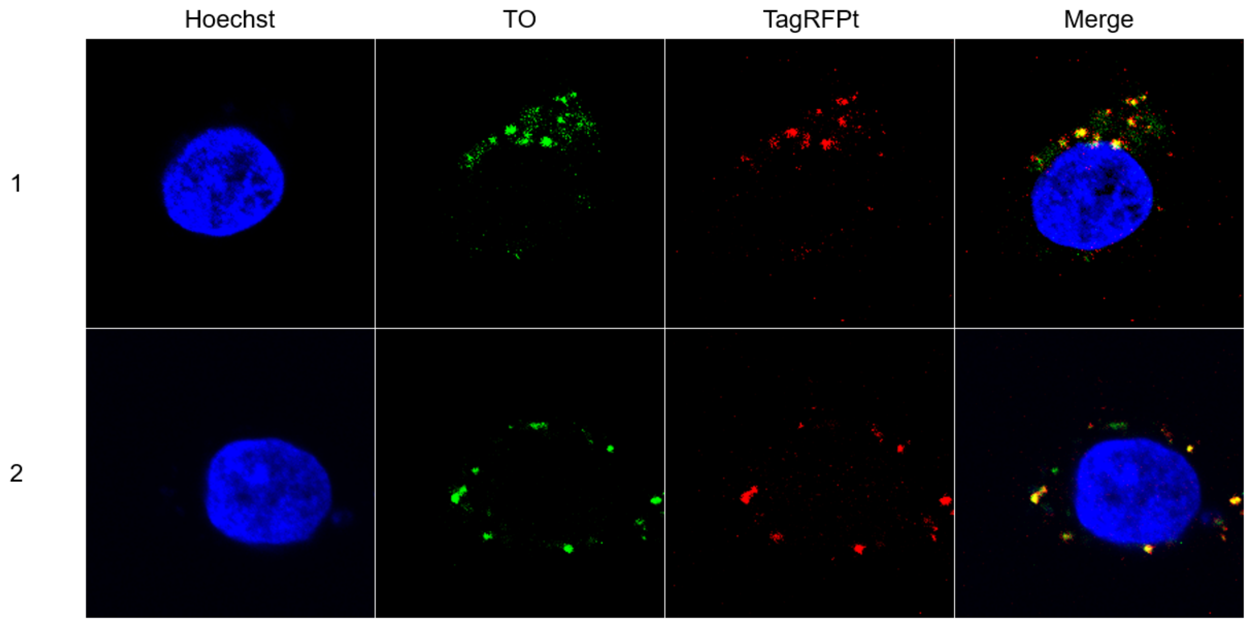


**Figure S3.1.** Colocalization of bPNA-TO and MCP-TagRFPt 2 hours after treatment with TO-bPNA. HEK-293T cells were transfected with the plasmids encoding MS2U4-tRNA and MCP-TagRFPt by Lipofectamine 3000. 1 μM K2M-Ala-K2M-TO was added for incubation at 37°C for 2 hours before imaging. Rows 1 and 2 were from two separate treatments; Row 1 is shown in manuscript Figure 5.

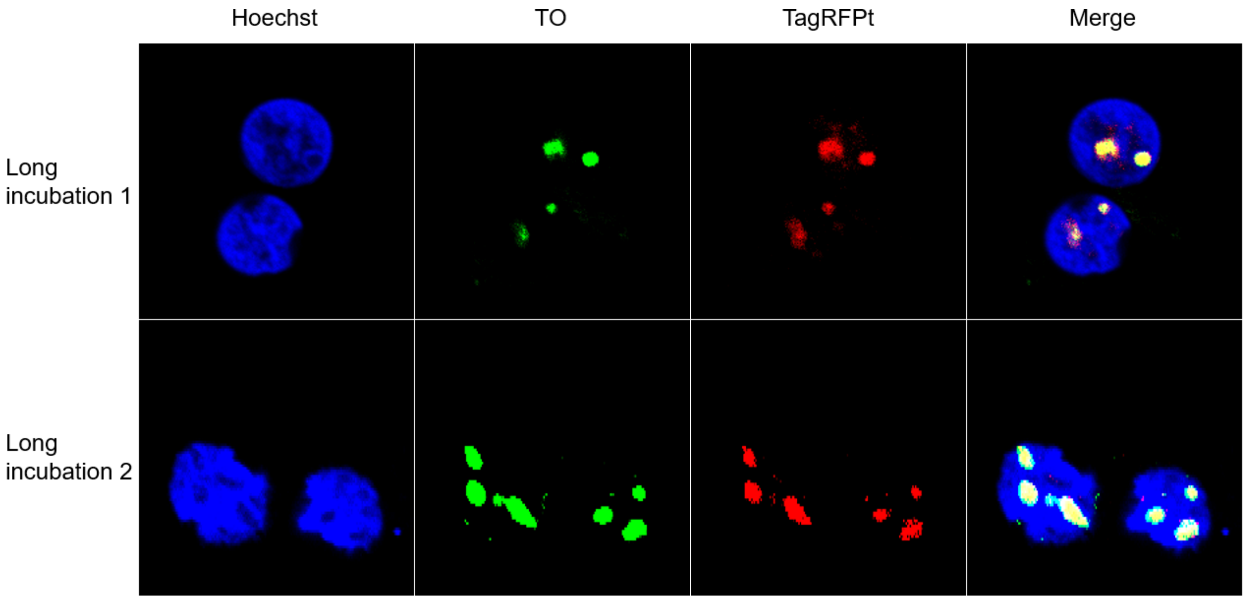


**Figure S3.2** Colocalization of bPNA-TO and MCP-TagRFPt 8 hr after treatment with TO-bPNA. HEK-293T cells were treated with MS2U4-tRNA, MCP-TagRFPt and bPNA-TO. Rows 1 and 2 were from two separate treatments; Row 1 is shown in manuscript Figure 5.

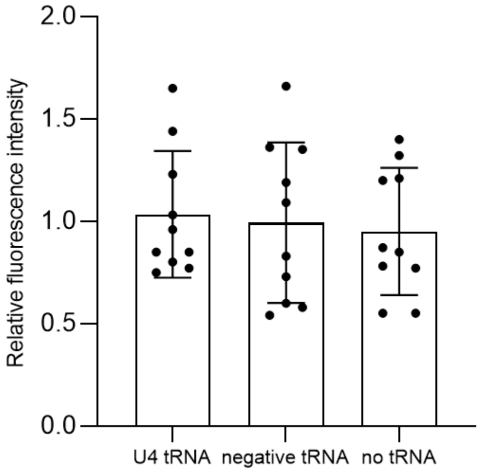


**Figure S3.3.** Cellular fluorescence intensity following treatment with Cy5-bPNA in place of TO-bPNA. HEK-293T cells were transfected as previously described with U4-tRNA, negative-tRNA or no tRNA. The relative intensities were normalized. 10 areas were selected for quantification for each type of treatment.

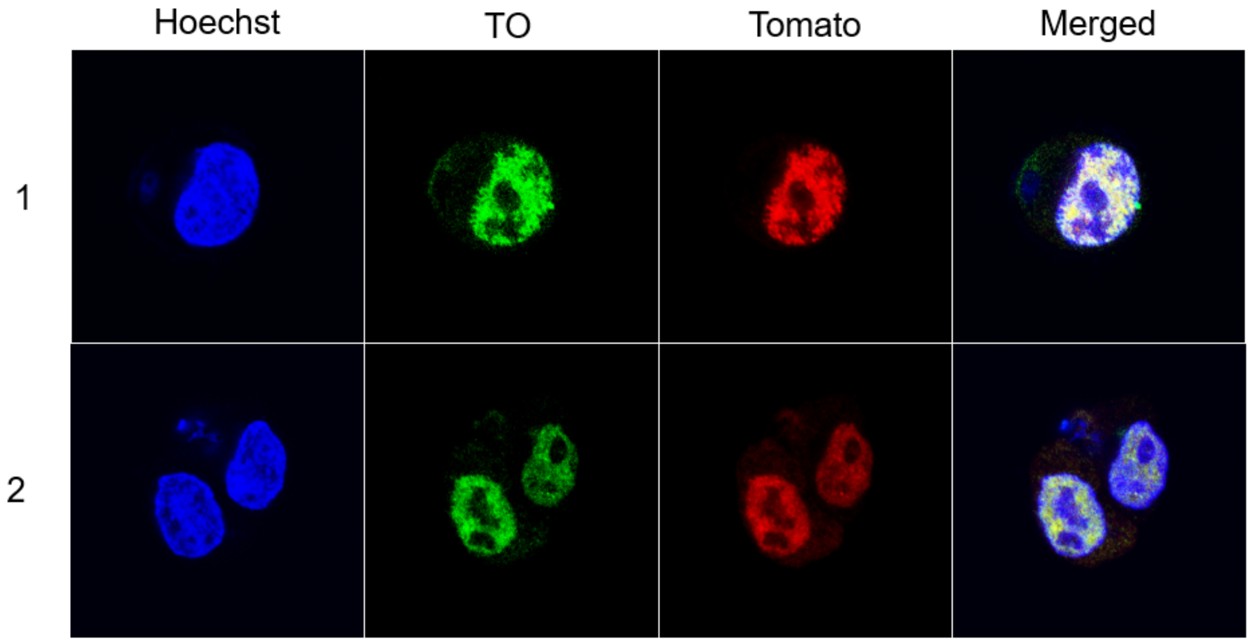


**Figure S3.4.** Additional replicate showing nuclear colocalization of TO-bPNA and TDP43-tdTomato fluorescence when co-expressed with U4-$(GU)_8$ in HEK-293T cells. Row 2 is shown in Figure 6, manuscript.

### S4. In vitro fluorogenic binding of TO-bPNA with RNAs.

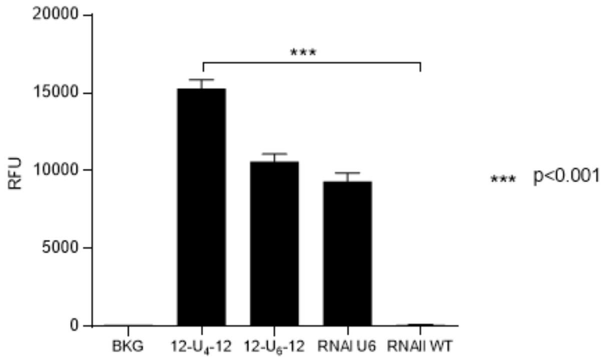


**Figure S4.1.** Fluorescence turn-on of TO-$K^{2M}$-Ala-$K^{2M}$ with RNAs and P-value as indicated.

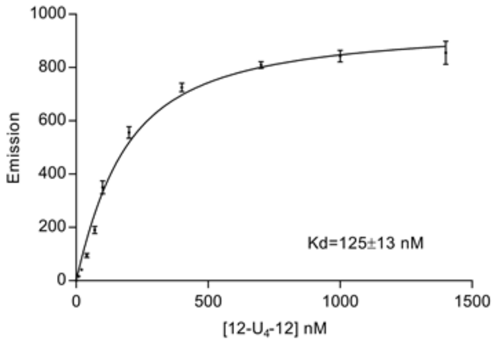


**Figure S4.2.** Fluorescence activation (apparent $K_d$) of TO-K2M-Ala-K2M with 12-$U_4$-12 RNA. Concentration of bPNA is 100 nM. Experimental conditions and fitting as described (S2.3).

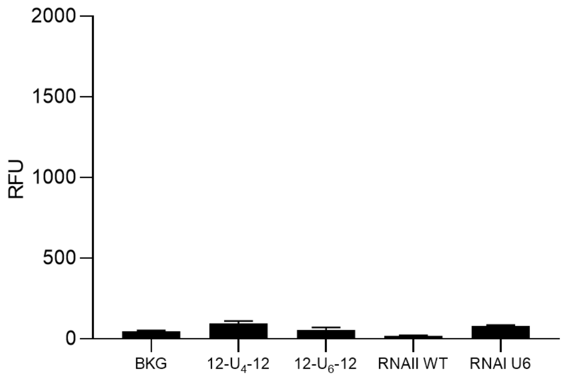


**Figure S4.4.** Fluorescence of DFHBI-K2M-Ala-K2M upon treatment with RNAs indicated. No fluorescence activation was observed.

### S5. Synthetic procedures

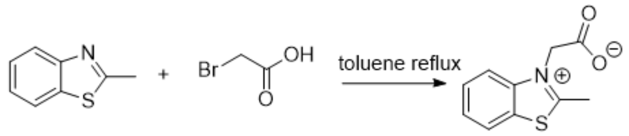


**Scheme S5.1.** Synthesis of 2-(2-Methylbenzo[d]thiazol-3-ium-3-yl)acetate (**1c**).

**2-(2-Methylbenzo[d]thiazol-3-ium-3-yl)acetate (2c).** The synthesis procedure was adapted from the reported procedure.[14] A mixture of 2-methylbenzothiazole (**1a**, 4.78 g, 32 mmol, 1.0 eq.) and bromoacetic acid (**1b**, 6.5 g, 47 mmol, 1.5 eq.) in toluene (100 ml) was heated at reflux overnight. After cooling to room temperature, the resulting mixture was filtered and the solid was washed with toluene (3 x 5 ml) to afford the product (**1c**, 6.5 g, 98%) after drying.

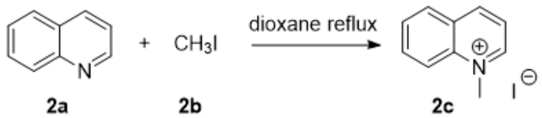


**Scheme S5.2.** Synthesis of 1-methylquinolin-1-ium iodide (**2c**).

**1-methylquinolin-1-ium iodide (2c).** The synthesis procedure was adapted from the reported procedure.[14] A mixture of quinoline (**2a**, 2 g, 15.5 mmol, 1.0 eq.) and iodomethane (**2b**, 2 ml, 32.2 mmol, 2.1 eq.) in 1,4-dioxane (150 ml) was heated at reflux for 1 hr. After cooling to room temperature, the resulting mixture was filtered and the solid was washed with diethyl ether (3 x 5 ml) and hexanes (3 x 5 ml), affording the product (**2c**, 4 g, 95%) after drying.

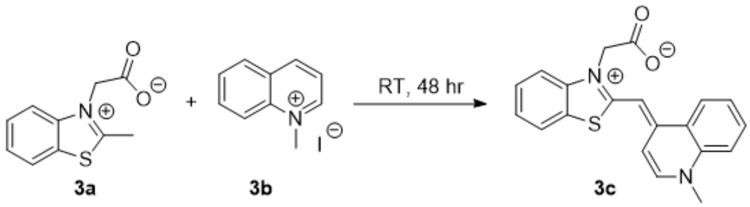


**Scheme S5.3.** Synthesis of TO-acetate (**3c**).

**TO-acetate (3c).** The synthesis procedure was adapted from the reported procedure.[14] 2-(2-Methylbenzo[d]thiazol-3-ium-3-yl)acetate (**3a**, 2.14 g, 10.4 mmol, 1.4 eq.), 1-methylquinolin-1-ium iodide (**3b**, 2 g, 7.4 mmol, 1 eq.) and triethylamine (2.24 g, 22.2 mmol, 3 eq.) were added in a round bottom flask with 30 ml dichloromethane. The mixture was stirred at room temperature for 48 hr. After 48 hr, the mixture was concentrated and 16 ml ethanol and 4 ml diethyl ether were added. The resulting mixture was stirred at 4 °C overnight. The resulting precipitate was collected by filtration and washed with diethyl ether (3 x 8 ml). The solid was then dissolved in 12 ml methanol with 3 ml water and was stirred overnight at 4 °C. The resulting precipitate was collected by filtration and washed with water (3 x 8 ml). Then the solid was resuspended in acetone and stirred at room temperature for 1 hr. The final product was obtained by filtration and was washed with acetone (3 x 8 ml) and dried in vacuum to afford TO-acetate as red solid (**3c**, 160 mg, 6.4%). **$^1$H NMR** (DMSO-d6): 8.54 (1H, d); 8.48 (1H, d); 7.98 (1H, d); 7.94 (1H, d); 7.86 (1H, t); 7.64 (1H, d); 7.55 (2H, m); 7.38 (1H, t); 7.18 (1H, d); 6.77 (1H, s); 5.09 (2H, s); 4.10 (3H, s). **HRMS** (ESI): Mass calculated: [M+H$^+$]=349.1005; Mass found: [M+H$^+$]=349.1004

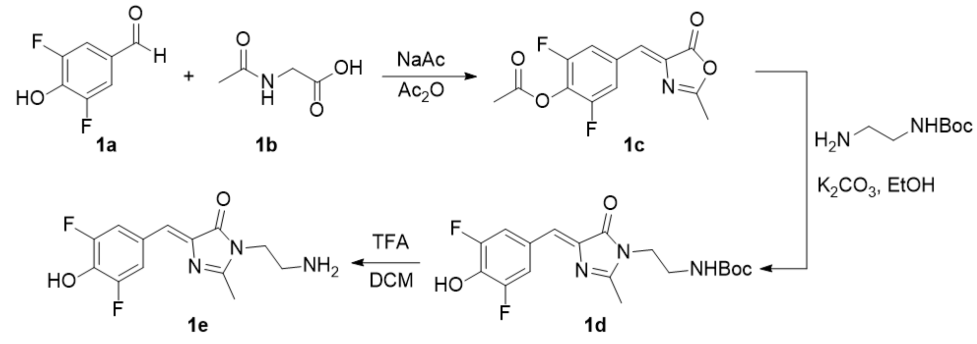

**Scheme S5.4.** Synthesis of (Z)-2,6-difluoro-4-((2-methyl-5-oxooxazol-4(5H)-ylidene)methyl)phenyl acetate.

**(Z)-2,6-difluoro-4-((2-methyl-5-oxooxazol-4(5H)-ylidene)methyl)phenyl acetate (1c).** The synthesis was adjusted from a reported procedure.[15] N-Acetylglycine (**1b**, 0.22 g, 1.9 mmol), anhydrous sodium acetate (0.156 g, 1.9 mmol), 4-hydroxy-3,5- difluorobenzaldehyde (**1a**, 0.3 g, 1.9 mmol), and acetic anhydride (0.72 ml) were stirred at 100 ºC for 2 h. Then the reaction was cooled to room temperature and 3 ml ethanol was added to the mixture. The mixture was stirred at 4 ºC overnight. The resulting solid was collected by filtration, washed with cold ethanol, hot water, hexanes and dried to afford 0.32 g (60%) of product as a yellow solid.

**DFHBI-ethylenediamine (1e).** The synthesis was adjusted from a reported procedure.[15] (Z)-2,6-difluoro-4-((2-methyl-5-oxooxazol-4(5H)-ylidene)methyl)phenyl acetate (**1c**, 100 mg, 0.36 mmol), Boc-ethylenediamine (110 mg, 0.69 mmol) and potassium carbonate (0.13 g) were added to 2 ml ethanol and refluxed for 4 hr. The mixture was cooled to room temperature and the solvent was removed in vacuum. The residue was redissolved in acetate buffer (pH=3.0) and ethyl acetate (1:1 mixture). The organic layer was collected and the solvent was removed in vacuum. The residue was purified by column (DCM:MeOH=10:1), yielding 104 mg (76%) product (**1d**) as yellow solid. The product was added to TFA:DCM=1:1 for Boc deprotection.

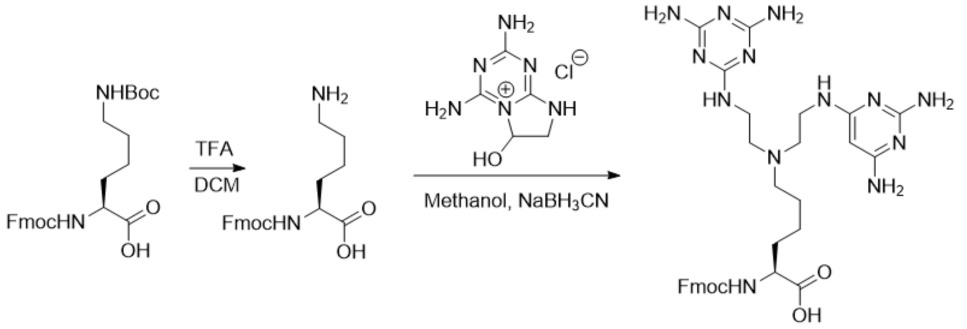

**Scheme S5.5.** Synthesis of Fmoc-$K^{2M}$-OH.

**Fmoc-$K^{2M}$-OH.** The procedure was the same as previously reported.[16] Fmoc-Lys(Boc)-OH (10 g, 21 mmol) was dissolved in 100 mL dichloromethane and 50 mL of trifluoroacetic acid was added. The reaction was stirred for 1 h and the solution was condensed to syrup under a stream of $N_2$. Dichloromethane (50 mL) was added to dissolve the syrup and was removed by a stream of $N_2$. The syrup was dissolved in 200 mL methanol and the pH was adjusted to 5 with solid $NaHCO_3$ . Melamine aldehyde (Hemiaminal form, 9.5 g, 46.2 mmol) and $NaBH_3CN$ (2.93 g, 46.2 mmol) was aliquoted into 4 portions, respectively. To the reaction solution, 2 portions of melamine aldehyde were added and were stirred and incubated at 50 ºC for 30 min. Then the reaction was taken out to cool to room temperature and 1 portion of $NaBH_3CN$ was added. The reaction was stirred at room temperature for another 30 min. Then 1 portion of aldehyde was added, incubated at 50 ºC for 30 min and 1 portion of $NaBH_3CN$ was added and incubated at room temperature for 30 min. These addition of aldehyde and reductant steps were repeated until all 4 portions of aldehyde and 3 portions of $NaBH_3CN$ were added. The last portion of $NaBH_3CN$ was added to the reaction and stirred for 30 min at room temperature. The reaction was monitored by HPLC. The remaining monoadduct (Fmoc-K1M-OH) was reacted by adding 2.3 g of melamine aldehyde, incubating at 50 ºC for 30 min and 0.72 g $NaBH_3CN$ was added to finish the reaction.

Methanol was reduced to ~50 ml and was discarded after centrifugation, yielding white solid as crude product after drying. The reaction was quenched by adding 10 mL 1N hydrochloric acid and the solid was triturated. The hydrochloric acid was discarded after centrifugation. Acetone (30 mL) was added to the residue and the residue was triturated. The acetone was removed by centrifugation. The acetone wash was repeated 3 times and ethanol was added for trituration instead of acetone. After removing ethanol by centrifugation, the product (~11 g, 78%) was obtained as white solid.

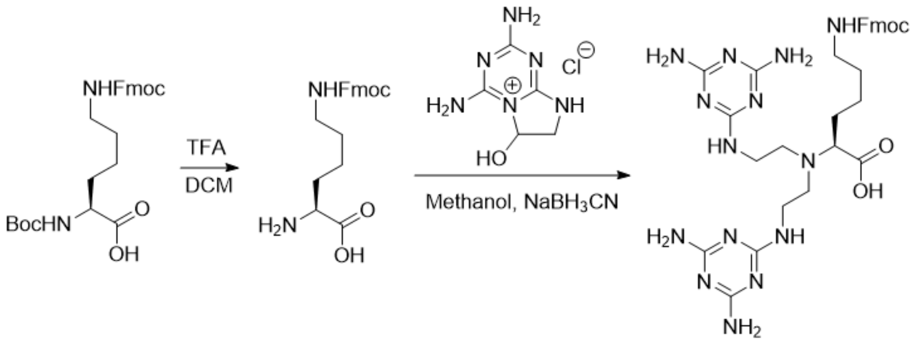


**Scheme S5.6.** Synthesis of Fmoc-α$K^{2M}$-OH.

**Fmoc-α$K^{2M}$-OH.** The procedure was the same as previously reported.[17] Boc-Lys(Fmoc)-OH (1 g, 2.1 mmol) was dissolved in 20 mL dichloromethane and 4 mL of trifluoroacetic acid was added. The reaction was stirred for 1 h and the solution was condensed to syrup under a stream of $N_2$. Dichloromethane (10 mL) was added to dissolve the syrup and was removed by a stream of $N_2$. The syrup was dissolved in 30 mL methanol and the pH was adjusted to 5 with solid $NaHCO_3$. Melamine aldehyde (0.95 g, 4.62 mmol) and $NaBH_3CN$ (0.293 g, 4.62 mmol) was aliquoted into 4 portions, respectively. To the reaction solution, 2 portions of melamine aldehyde were added and incubated at 50˚C for 30 min with stirring. Then the reaction was taken out to cool to room temperature and 1 portion of $NaBH_3CN$ was added. The reaction was stirred at room temperature for another 40 min. Then 1 portion of aldehyde was added, incubated at 50˚C for 30 min and 1 portion of $NaBH_3CN$ was added and incubated at room temperature for 40 min. These additions of aldehyde and reductant steps were repeated until all 4 portions of aldehyde and 3 portions of $NaBH_3CN$ were added. The last portion of $NaBH_3CN$ was added to the reaction and stirred for 40 min at room temperature. The reaction was monitored by HPLC. The remaining monoadduct (Fmoc-α$K^M$-OH) was reacted by adding 0.25 g of melamine aldehyde, incubating at 50˚C for 30 min and 0.078 g $NaBH_3CN$ was added to finish the reaction. Methanol was reduced to 5 ml and was discarded after centrifugation, yielding pale yellow solid as crude product after drying. The reaction was quenched by adding 2 mL 1N hydrochloric acid and the solid was triturated. The hydrochloric acid was discarded after centrifugation. Acetone (5 mL) was added to the residue and the residue was triturated. The acetone was removed by centrifugation. The acetone wash was repeated 3 times and ethanol was added for trituration instead of acetone. After removing ethanol by centrifugation, the product (0.85 g, 60%) was obtained as pale yellow solid. **$^1$H NMR** (400 MHz, DMSO-d6): 7.91 (2H, d); 7.79 (10H, d); 7.66 (2H, d); 7.41 (2H, t); 7.32 (2H, t); 7.25 (1H, t); 4.28 (2H, d); 4.20 (1H, t); 3.33 (4H, q); 3.28 (1H, m); 2.95 (4H, q); 2.76 (2H, q); 1.62 (2H, q); 1.15-1.50 (4H, m). **$^{13}$C NMR** (100 MHz, DMSO): 174.3; 170.7; 166.3; 154.5; 144.3; 141.1; 128.0; 127.5; 125.5; 120.5; 80.1; 79.6; 65.5; 50.6; 49.0; 47.2; 29.3; 23.6; 21.9. **HRMS** (ESI): calculated for [M+H]=673.3430, [M+2H]=337.1751, found [M+H]=673.3425, [M+2H]=337.1756.

**S6. Solid phase peptide synthesis**

Peptide synthesis was performed manually using Rink Amide resin (100-200 mesh, loading 0.3 mmol/g) employing standard Fmoc chemistry. With 150 mg resin, 0.25 M of Amino acids were coupled with 0.25 M of PyAOP and 0.25 M DIPEA in 2 ml NMP. Fluorogenic dyes were coupled using three equivalents of dye, 3.3 equivalents of HBTU, and 3.3 equivalents of DIPEA in 2 ml DMF. Fmoc cleavage was performed with 2 mL of piperidine:NMP (1:1) with 3% DBU. Dye and coupling reagents were allowed to react for 15 min before addition to resin. For TO peptides, TO- acetate was coupled to the peptide N-terminus directly. For DFHBI peptide, the peptide was reacted with 3 equivalents of succinic anhydride, and 3 equivalents of DIPEA for 15 min,washed and DFHBI-$NH_2$ was coupled to the N-terminus of the peptide. Peptides were cleaved from the solid support using 95% trifluoroacetic acid (TFA) and 5% $H_2O$ for 2 h. Cold diethyl ether ($Et_2O$) was added to precipitate the peptide and the crude pellet was washed with cold $Et_2O$ two times and dried over vacuum. Crude peptides were then dissolved in solvent A and purified by HPLC on a semi-prep $C_{18}$ reversed phase column at 8 mL/min. The UV detector was set at 238 nm. The purified peptides were lyophilized to dryness. The identity of peptide was checked by MALDI-TOF and purity checked by analytical HPLC on a $C_{18}$ column. (solvent A=0.1%TFA in water, solvent B=0.07% TFA in 90% acetonitrile, 10% water).

### S7 Compound characterization.

- **TO-acetate**

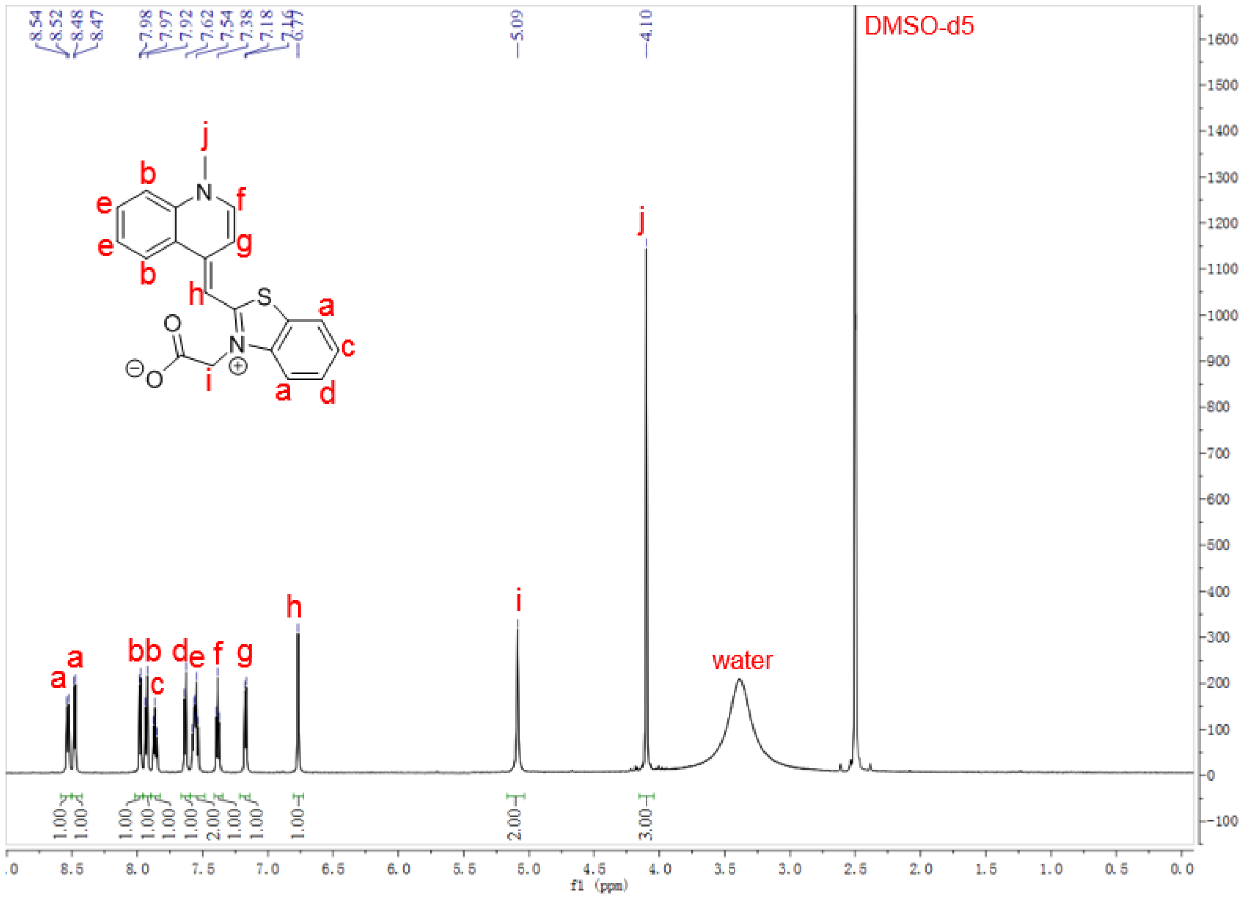


**Figure S7.1.** $^1$H NMR of TO-acetate. (DMSO-d6)

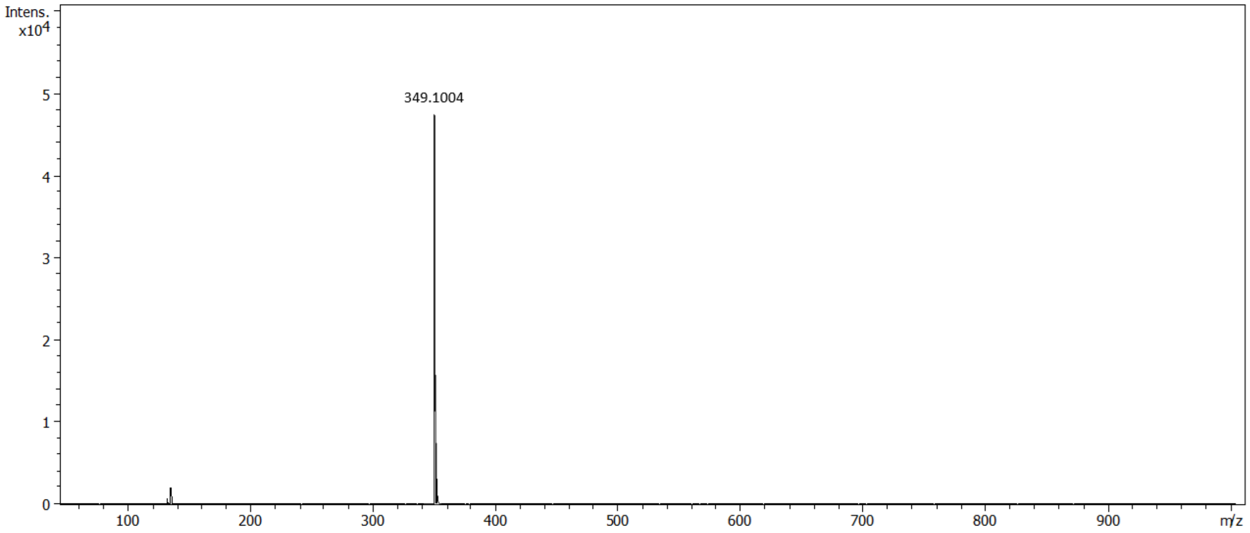

**Figure S7.2.** ESI of TO-acetate. Mass expt: [M+H$^+$]=349.1004; Mass calc: [M+H$^+$]=349.1005.

- **DFHBI-ethylenediamine-Boc**

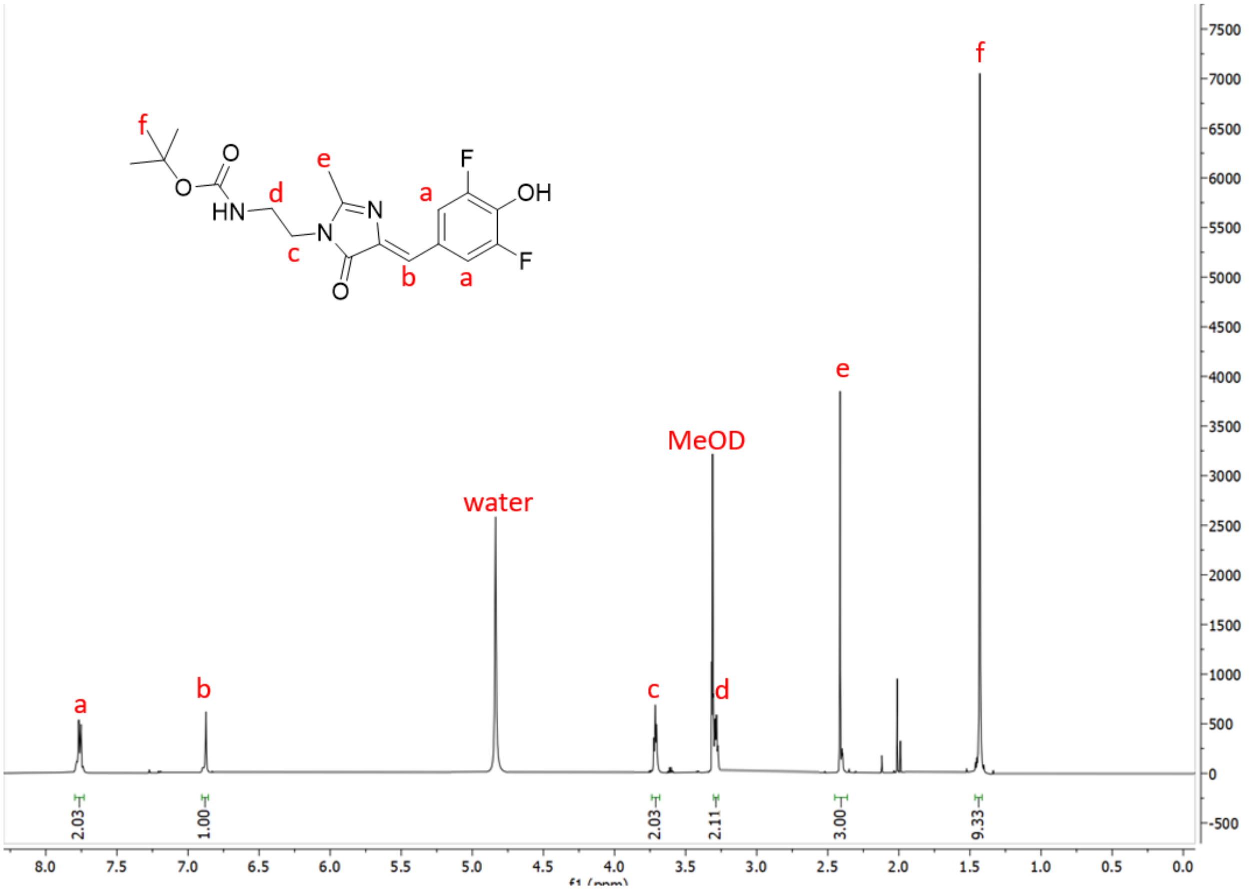


**Figure S7.3.** $^{1}$H NMR of DFHBI-ethylenediamine-Boc. (DMSO-d6)

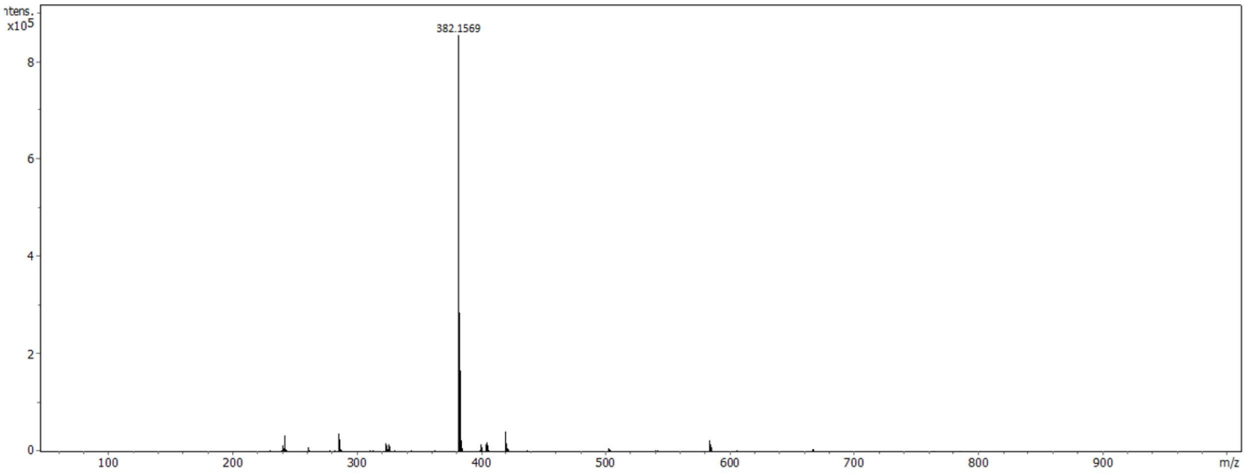


**Figure S7.4.** ESI of DFHBI-ethylenediamine-Boc. Mass expt: $[M+H^{+}]$=382.1569; Mass calc: $[M+H^{+}]$=382.1573.

- **$K^{2M}$-$K^{TO}$-$K^{2M}$**

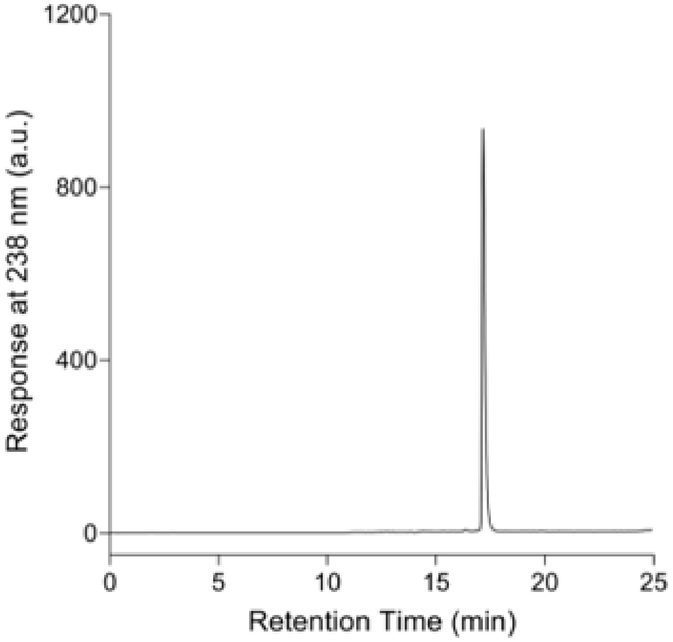


**Figure S7.5.** HPLC of $K^{2M}$-$K^{TO}$-$K^{2M}$. Gradient: 0-5 min: 0% B; 5-15 min: 0-40% B; 15-20 min: 40% B; 20-20.5 min: 40-100% B; 20.5-22.5 min: 100% B; 22.5-23 min: 100-0% B; 23-25 min: 0% B.

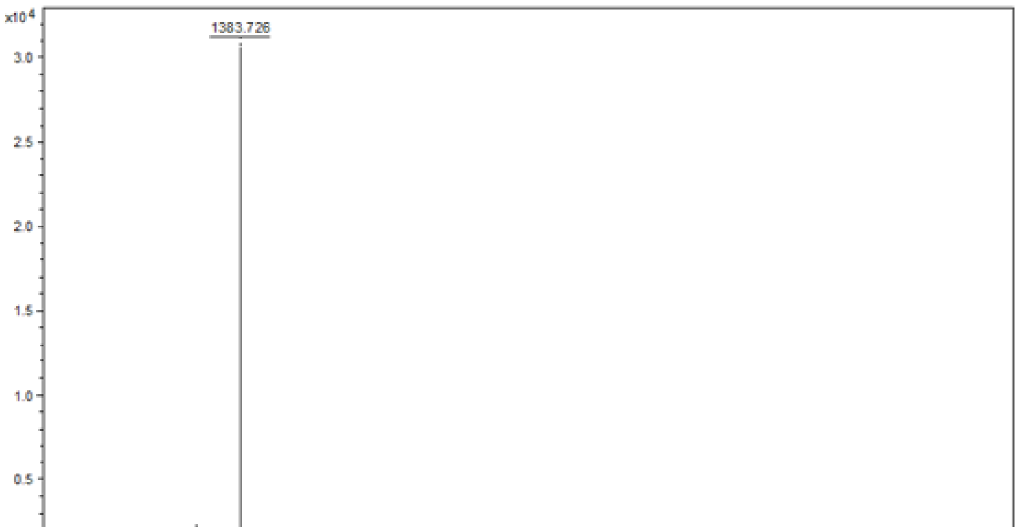


**Figure S7.6.** MALDI-TOF of Ac-$K^{2M}$-$K^{TO}$-$K^{2M}$. Mass calculated: $[M+H^+]$=1383.743; Mass found: $[M+H^+]$=1383.726.

- **TO-βAla-K$^{2M}$-Ala-K$^{2M}$**

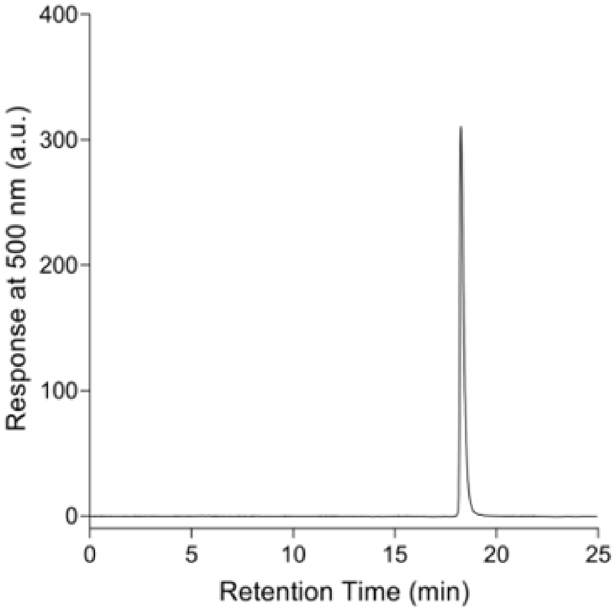


**Figure S7.7.** HPLC of TO-βAla-K$^{2M}$-Ala-K$^{2M}$. Gradient: 0-5 min: 0% B; 5-15 min: 0-40% B; 15-20 min: 40% B; 20-20.5 min: 40-100% B; 20.5-22.5 min: 100% B; 22.5-23 min: 100-0% B; 23-25 min: 0% B.

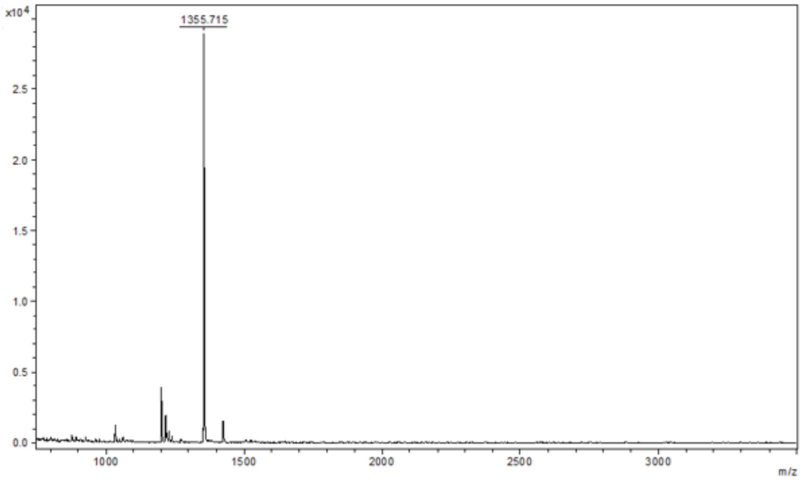


**Figure S7.8.** MALDI-TOF of TO-βAla-K$^{2M}$-Ala-K$^{2M}$. Mass calculated: [M+H$^+$]=1355.712; Mass found: [M+H$^+$]=1355.715.

- **TO-βAla-K$^{2M}$-Ile-K$^{2M}$**

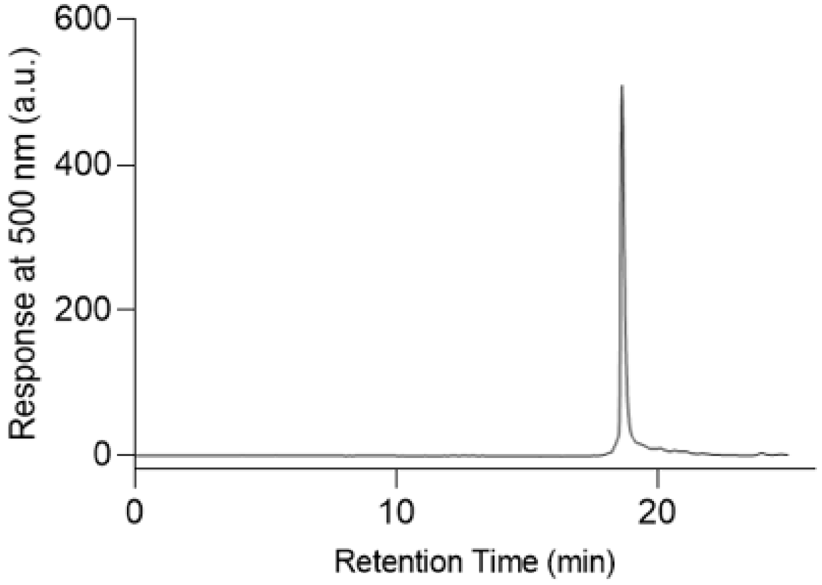


**Figure S7.9.** HPLC of TO-βAla-K$^{2M}$-Ile-K$^{2M}$. Gradient: 0-5 min: 0% B; 5-15 min: 0-40% B; 15-20 min: 40% B; 20-20.5 min: 40-100% B; 20.5-22.5 min: 100% B; 22.5-23 min: 100-0% B; 23-25 min: 0% B.

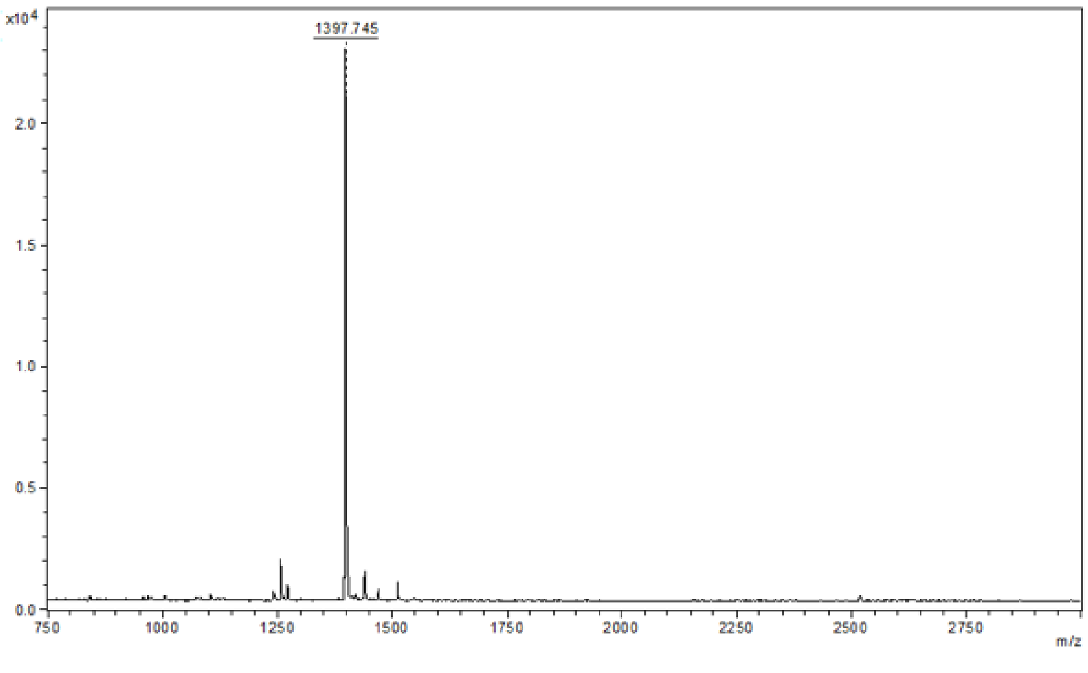


**Figure S7.10.** MALDI-TOF of TO-βAla-K$^{2M}$-Ile-K$^{2M}$. Mass calculated: [M+H$^+$]=1397.759; Mass found: [M+H$^+$]=1397.745.

- **TO-iso($K^{2M}$)$_2$**

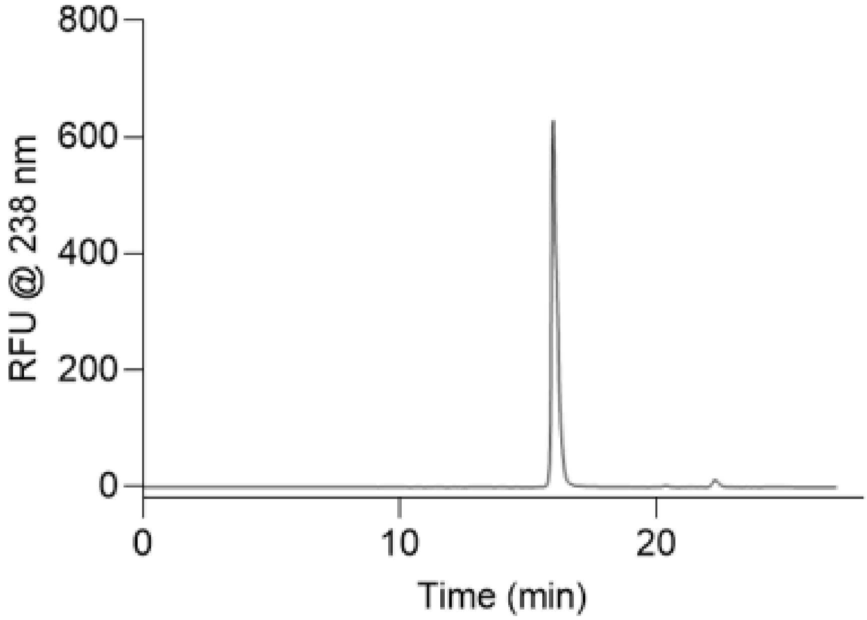


**Figure S7.11.** HPLC of TO-iso($K^{2M}$)$_2$. Gradient: 0-5 min: 0% B; 5-15 min: 0-40% B; 15-20 min: 40% B; 20-20.5 min: 40-100% B; 20.5-22.5 min: 100% B; 22.5-23 min: 100-0% B; 23-25 min: 0% B.

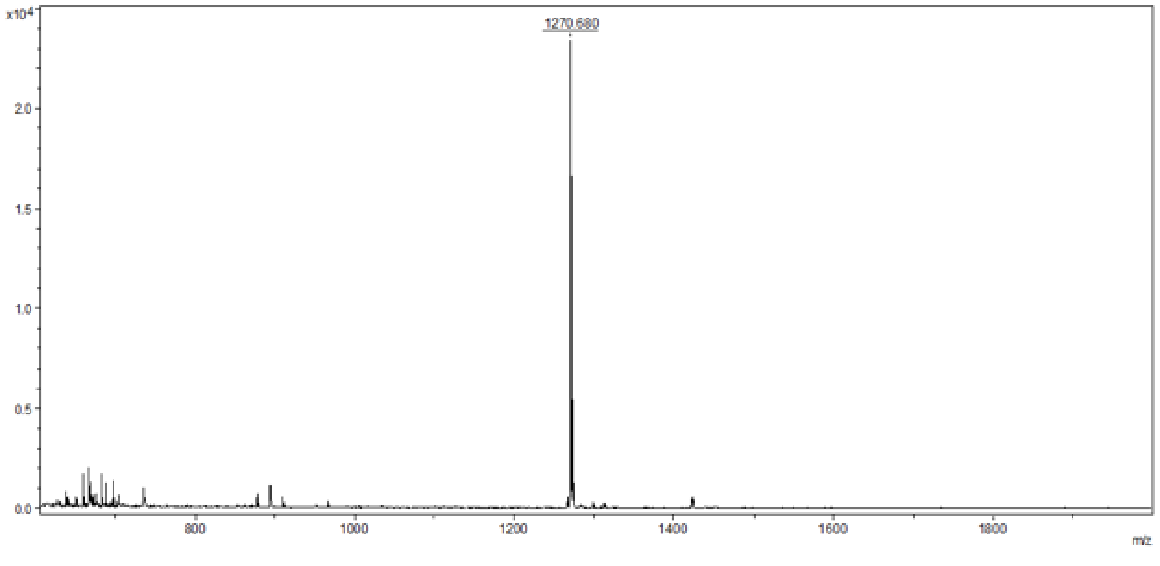


**Figure S7.12.** MALDI-TOF of TO-iso($K^{2M}$)$_2$. Mass calculated: [$M+H^+$]=1270.659; Mass found: [$M+H^+$]=1270.680.

- **DFHBI-K$^{2M}$-Ala-K$^{2M}$**

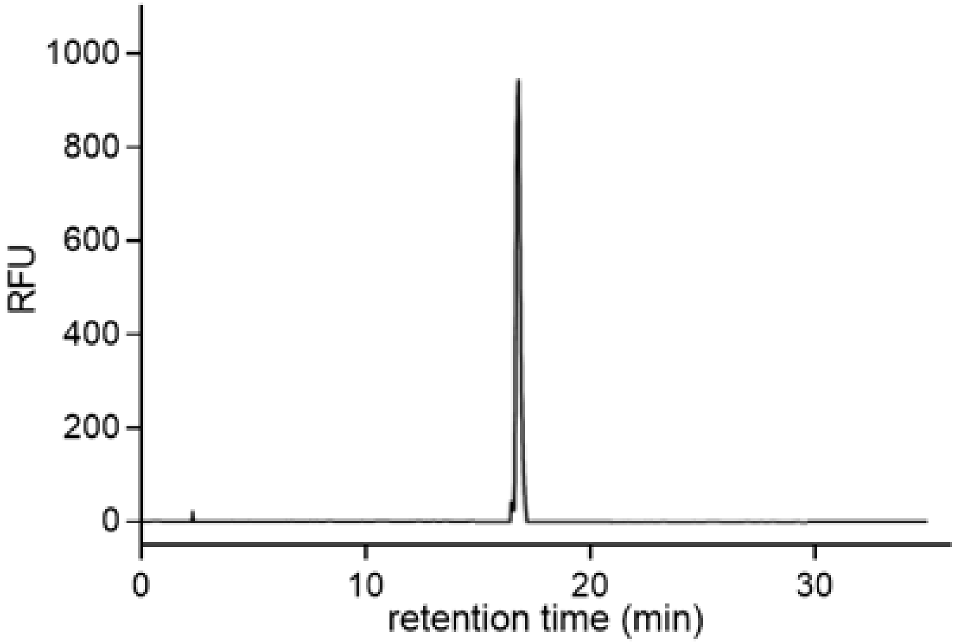


**Figure S7.13.** HPLC of DFHBI-K$^{2M}$-Ala-K$^{2M}$. Gradient: 0-5 min: 0% B; 5-15 min: 0-40% B; 15-20 min: 40% B; 20-20.5 min: 40-100% B; 20.5-22.5 min: 100% B; 22.5-23 min: 100-0% B; 23-25 min: 0% B.

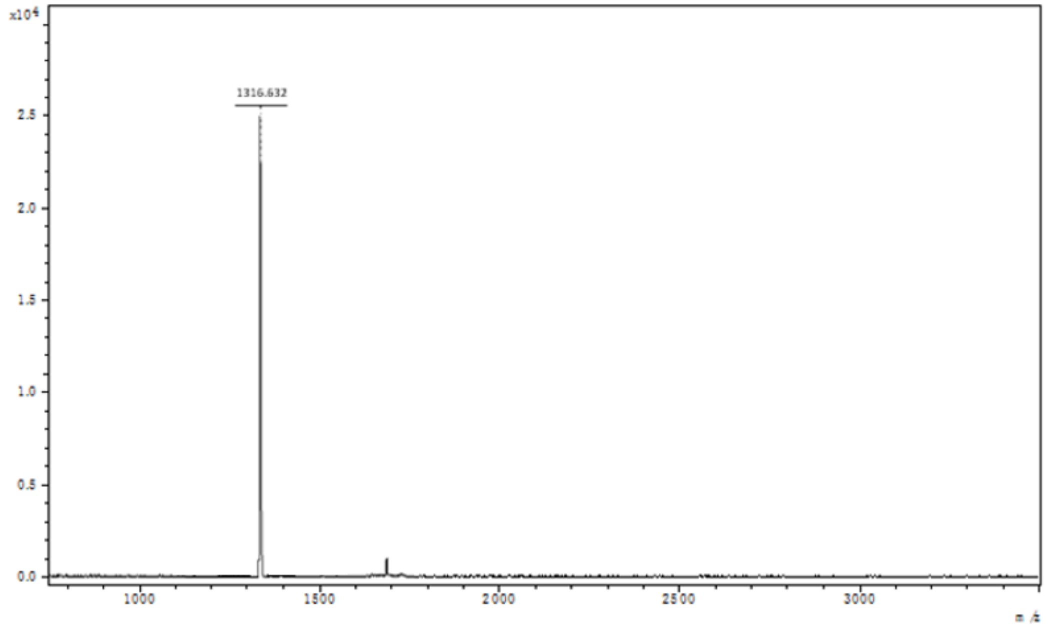


**Figure S7.14.** MALDI-TOF of DFHBI-K$^{2M}$-Ala-K$^{2M}$. Mass calculated: [M+H$^+$]=1316.688; Mass found: [M+H$^+$]=1316.632.
